# Mechanical coupling narrows an epithelial colony before it reaches a bottleneck

**DOI:** 10.64898/2026.09.12.751083

**Authors:** Sun-Min Yu, Yoon-Kyoung Cho, Steve Granick

## Abstract

A colony migrating into a passage narrower than itself but wider than individuals must adapt its density distribution to pass. Powders, colloids and crowds driven through such an opening jam, since contact interactions provide no pathway for constituents to respond to the constriction before reaching it. Exploring whether a cohesive living tissue can reorganize in advance, we image MDCK epithelial monolayers migrating through microchannels in which a wide channel narrows, and locate response at the taper mouth, a transition zone in which we identify upstream responses. At locations where the local channel width is still unchanged, the monolayer detaches from the side walls and reduces its lateral extent. Particle image velocimetry shows that this is not passive compression; the axial velocity alternates aperiodically in sign, velocity divergence is positive in the transition zone, and tracking the positions of cell nuclei confirms that cells leave the crowded inlet region in both directions. Fluorescence imaging shows a belt of co-enriched E-cadherin, F-actin, and myosin-II at the edges that detach before the inlet, which laser ablation shows to be under elevated tension but dissolves once the tissue passes into the narrow inlet; β-catenin is not co-enriched, indicating that junctions differ in composition in the core and at the edge of colony. This advance proceeds over hours, far more slowly than the viscoelastic reorganization reported for MDCK monolayers, and switches in sign repeatedly, which a linear viscoelastic sheet under sustained constraint cannot do. E-cadherin knockdown abolishes narrowing, flow reversal, density redistribution, and belt assembly.

**The bigger picture:** Cell collectives squeeze through narrow passages during development, wound repair, and cancer invasion, often meeting gaps wider than one cell but narrower than the group. Powders and crowds of people jam at such openings because no constituent learns of the obstruction before arriving. Here we show that epithelial cell colonies do not. When the leading edge enters a narrowing channel, cells far behind it redistribute themselves and pull the sheet off the side walls, so the group is already narrower on arrival. Removing the adhesion protein linking neighboring cells eliminates this response. Mechanical coupling therefore carries information backward through a tissue, letting following cells reshape the colony before it reaches the geometrical constraint.

## Introduction

Navigating confined spaces enables multicellular structures to respond to external stimuli (1-3). Paradoxically, it is also an essential process to achieve cancer cell metastases (4, 5), which raises the possibility that a small number of generic migration scenarios are shared across these processes. Tissue mechanics is one route by which collective features are controlled (6, 7), but the demands of confinement differ according to its severity.

In extreme confinement, where passages are narrower than individual cells, cell nuclei are deformed, activating mechanosensitive pathways and altering gene expression (1, 8); at the opposite limit, uniform straight channels impose constant geometric constraints. Considerable research has established how confinement in this regime enhances directional persistence, generates spontaneous flow, and organizes leader-follower dynamics (9-11). In these systems, most studies report that information flows from leaders to followers (9, 12-15). The resulting mechanical waves have been characterized during unconstrained expansion (12) or in straight channels (9, 11, 16), rather than in response to geometric obstacles. Here, we are concerned with a less-studied regime between these limits: passages that are narrower than a migrating colony but wider than individual cells, such that the collective must adapt its shape, density distribution, and internal organization to transit the constriction, analogous to the discharge of a crowd through a doorway or of grains through a silo orifice.

Outside biology this is the bottleneck problem, whose generic outcome is clogging. Powders, colloidal suspensions, sheep herds and even human crowds form transient arches at the opening when they discharge through an aperture (17). The throughput of pedestrian crowds grows with exit width but congestion can develop even when the incoming flux is below its nominal capacity (18), and at high density the flow switches from smooth to stop-and-go and then disordered (19). Interestingly, information is transmitted upstream: speed perturbations in polarized crowds can propagate as weakly damped backward-traveling waves (20), and confined crowds above a critical density develop collective oscillations (21). But the constituents interact only by contact repulsion, so upstream reorganization is limited to the consequences of steric contact. Moreover, adding attraction raises the clogging (22).

Cells in an epithelial colony differ fundamentally from cohesive grains in that they generate contractile stress and also change state in response to the stress that they experience. Epithelial sheets transmit mechanical forces over multi-cell length scales through E-cadherin-based adherent junctions, which form mechanically continuous connections across the tissue (4, 23, 24). Force transmission in cell monolayers is characterized by intercellular stress fluctuations that propagate across the sheet and orient local migration long-ranged (25). When cells at the migration front experience increased lateral confinement, the resulting changes in cytoskeletal tension could propagate backward through these junctions, potentially activating mechanosensitive responses in upstream cells that have not themselves met the constraint. We refer to this as retrograde mechanical coupling, using retrograde with respect to the direction of colony migration. Leader-follower models developed for straight channels treat the follower response as information propagating from the front (9, 11, 16) and do not address this case.

For experiments, we designed tapered microchannels that impose a controlled geometric transition on migrating MDCK epithelial monolayers. Combining particle image velocimetry, live fluorescence imaging, laser ablation, and pharmacological and genetic perturbations, we find that colonies exhibit a coordinated retrograde response: upstream cells narrow their collective width, dynamically redistribute cell density through oscillatory flow, and assemble a peripheral belt of co-enriched E-cadherin, F-actin, and myosin-II at the colony edges under elevated mechanical tension. β-catenin is not co-enriched at this belt, indicating that peripheral junctions differ in molecular composition from interior junctions. This reorganization begins before upstream cells reach the inlet and is reversed after the colony exits confinement. E-cadherin knockdown abolishes every component of this response. These findings identify retrograde mechanical coupling as the mode of colony-scale shape adaptation upstream of geometric constrictions.

## Results

### Terminology

We define Zones I, II and III (Fig. 1a) according to their position relative to the constriction inlet (9, 11, 16). Previous literature about cell migration has defined leader and follower identities in the context of tumor cell invasion and along the epithelial-mesenchymal transition continuum (26) but did not address geometrical constrictions that concern us here, so we avoid this language. Position is measured from a point 200 µm beyond the start of the narrowest channel, with Zone I defined as 0–200 µm, Zone II as 200–600 µm, and Zone III as 600–1000 µm, denoted by the horizontal lines in Fig. 1a.

**Figure 1.**
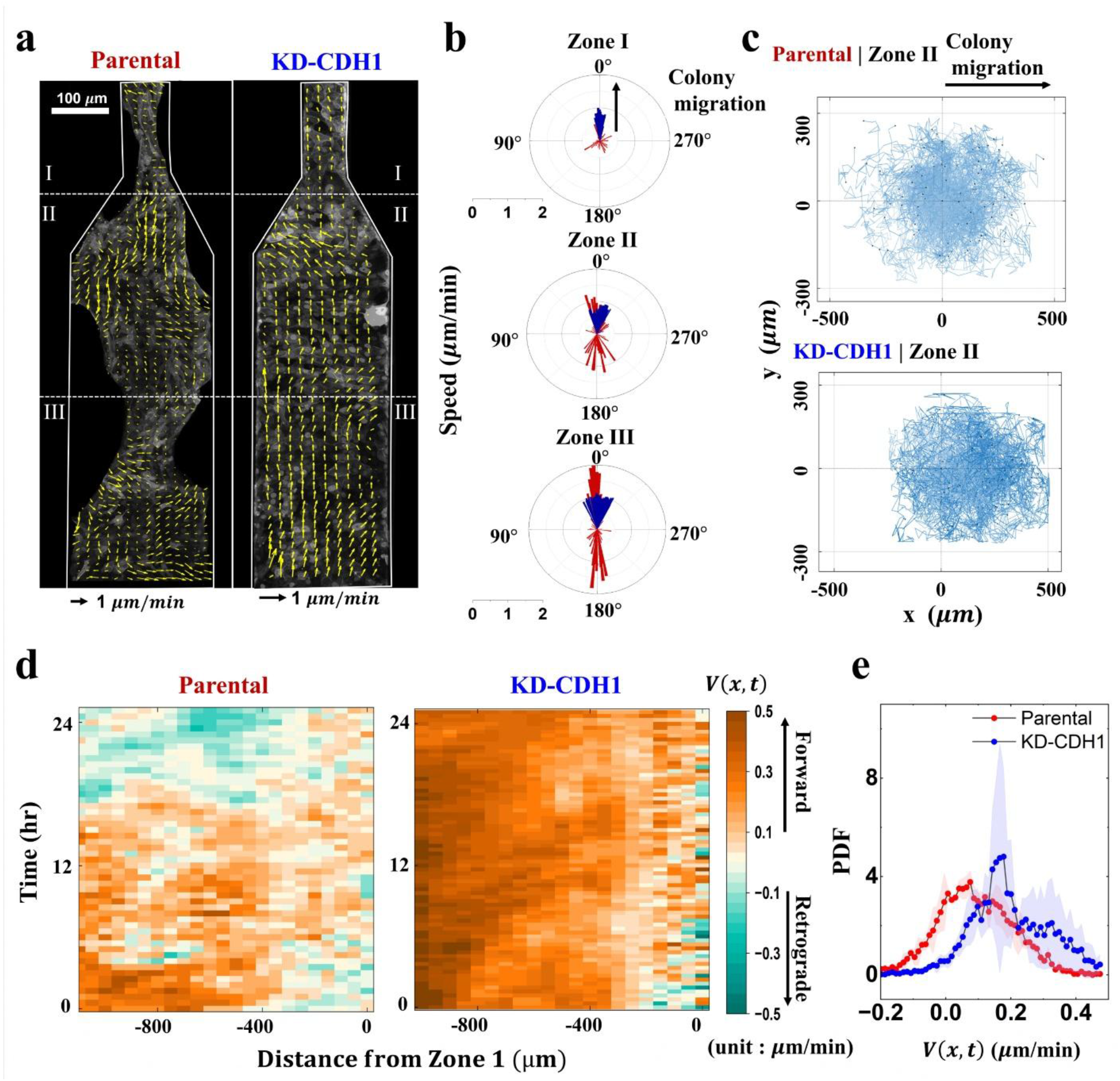
Parental colonies alternate forward and retrograde flow at a tapered inlet; KD-CDH1 colonies do not. (a) PIV velocity fields at 10 h for parental (left) and KD-CDH1 (right) colonies co-expressing H2B-mCherry (nuclei) and lifeact-mEGFP. Dashed lines mark demarcation between Zones I, II and III. (b) Polar distributions of velocity at 10 h in Zones I, II, and III. Black arrow shows direction of forward migration. (c) Angular distributions of displacement from individually tracked cell nuclei in Zone II, parental and KD-CDH1, evaluated over 30 and 20 min intervals, respectively. (d) Kymographs of the axial velocity, *V*(*x*,t), with x=0 located 200 µm into Zone I, averaged across width of the colony. (e) Probability density of *V*(*x*,t) for parental and KD-CDH1 colonies. N=3 independent experiments. Comparing parental colonies to those with KD-CDH1, we show that the mean and standard deviations of the fraction of retrograde velocity were 22±6.2% and 4±0.6%, respectively.

### Migrating colonies undergo collective reorganization at tapered inlets

Colonies of parental MDCK were compared to a cell line whose cadherin expression we knocked down (KD-CDH1) as described below. Monolayers co-expressing H2B-mCherry and Lifeact-mEGFP were imaged as they migrated through microchannels (Fig. S1, S2) with tapered inlets connecting a wide channel (*w* = 250 μm) to a narrow channel of width *w’* (Fig. S1b). Unless noted otherwise, experiments were performed in uncoated PDMS channels bonded to glass. For adhesion-control experiments, PDMS channels were coated with fibronectin (10 μg/ml) to increase cell–wall adhesion and test whether wall adhesion altered boundary detachment. Taper angle varied systematically (Fig. S2); subsequent experiments used *θ* = 30°. Migration was independent of channel height over the range 20 to 150 μm. The narrow channel width *w’* = 80 μm was selected from a screen of *w*’ values (20 to 150 μm) as the width producing maximal upstream area reduction and longest spatial correlation lengths.

Cell colonies initially occupied full channel width and migrated as a cohesive front. Upon reaching the tapered inlet, three coordinated responses emerged. First, leading cells accelerated and elongated along the migration axis, consistent with established single-cell responses to lateral confinement (1, 8). The cells behind began to narrow before following cells reached the inlet: 150 to 200 μm upstream of the constriction it detached from channel walls and the colony reduced its lateral extent, indicating reorganization in advance of direct encounter with the geometric constraint (Fig. 1a). Colony area in the wide channel decreased by 18 ± 12% over 24 h for parental cells (Fig. S3b). Third, particle image velocimetry (PIV) analysis showed that this narrowing was not passive compression: cells in Zones II and III migrated with both forward and backward velocity components (Fig. 1b-parental, Movie S1). Single-cell analysis of the polar velocity distributions of cell nuclei shows that individual cells in both parental and KD-CDH1 cell lines span all in-plane directions but skewed, for KD-CDH1 cells, more uniformly in the forward direction (Fig. 1c). The same appeared using other inlet geometries, including a gently curved channel (Fig. S3c-j, Movie S2), so it is not specific to this specific taper.

### Reorganization requires both intercellular adhesion and intracellular contractility

Using tetracycline-inducible shRNA, we reduced E-cadherin expression to 40 ± 1.5% of wild-type levels (Fig. S4, Fig. S5 a-b). At this expression level, cells formed morphologically normal monolayers with intact cell-cell contacts, yet KD-CDH1 colonies showed no area reduction over 24 h (Fig. S3b). Flow was predominantly unidirectional and forward, without upstream narrowing (Fig. 1b, 1c, Movie S3). By 20 h, parental cells carried both forward and backward velocity components at the inlet; KD-CDH1 cells retained narrow forward velocity distribution (Fig. 1b). Single-cell tracking gave the same result, with KD-CDH1 velocities concentrated in the forward direction (Fig. 1c).

To resolve motion along the net direction of migration, we projected the velocity field onto the x axis and averaged over y, giving *V*(*x, t*), which name as axial velocity. Kymographs of this quantity distinguish forward (*V*>0) from retrograde motion (*V*<0). KD-CDH1 colonies moved forward almost uniformly but parental colonies alternated aperiodically between forward and retrograde motion (Fig. 1d). The probability distribution of *V* shifted towards higher positive values upon KD-CDH1 knockdown (Fig. 1e). The mean and standard deviation value of the fraction of retrograde velocity (*V*<0) was 22±6.2% and 4±0.6% of parental and KD-CDH1 cells, respectively (Fig. 1e). The response was not specific to the tapered geometry: colonies entering a serrated inlet showed the same pattern of forward and retrograde motion, with nucleus displacement normal to the surface remaining below ± 2*µm* (Fig. S6), confirming that the retrograde signal reflects in-plane motion.

Reorganization was not prevented by increased wall adhesiveness nor on three-dimensional tissue organization. Fibronectin coating (10 μg/ml) delayed the onset of wall detachment by approximately 4 h but did not prevent reorganization in parental cells (Movie S4, Fig. S2a-b). KD-CDH1 cells did not respond to surface treatment, indicating that loss of mechanical connectivity, not altered cell-substrate adhesion, accounts for the failure of reorganization. Nuclei in both parental and KD-CDH1 colonies remained close to a single plane without out-of-plane structure (Fig. S7). Parental colonies fluctuated more in this z direction, most strongly at the colony boundary, yet the fluctuation remained within ± 2 *µm* throughout (Fig. S7c), so the colony can be treated as a monolayer and the reorganization as an in-plane process. This enhanced boundary fluctuation is consistent with the boundary contractility described below.

We next computed the two-dimensional divergence of the PIV velocity field, ∇·v, which localizes coordinated expansion and compression. In parental colonies, the distribution of the mean divergence in Zone II shifted toward positive values with a Gaussian center of 0.13± 0.05 *h*^−1^, compared with 0.05 ± 0.06 *h*^−1^in Zone I and near zero in the Zone III (Fig. S8b, Table S3). KD-CDH1 colonies did not show this regional separation, with the three zones indistinguishable (Fig. S8c, Table S3). Sustained positive mean divergence in a confluent sheet requires that cells leave the plane, that cell number density fall, or that individual cell area increase. The nuclei z positions exclude the first one (Fig. S7).

Together, parental colonies narrow their width and alternate flow direction upstream of the constriction, responding actively to geometric constraint rather than deforming passively. Disrupting E-cadherin-based force transmission eliminates both responses. The sections that follow test three components of this behavior: the molecular and mechanical requirements for collective reorganization, the redistribution of cell density by oscillatory flow, and the structural basis of retrograde force transmission. These depend on E-cadherin, as knockdown abolishes reorganization, eliminates oscillational flow, and prevents the formation of a peripheral mechanical belt. This shared dependence motivates the working model that a peripheral belt provides mechanical infrastructure for retrograde force transmission, oscillational flow is the dynamic manifestation of that transmission, and colony narrowing is the functional outcome.

### Pharmacological perturbations confirm requirements for contractility and adhesion

If retrograde reorganization requires both intracellular force generation and intercellular force transmission, then disrupting either component should impair reorganization. Pharmacological perturbations confirmed this prediction (Fig. S5 c-e; full description in Table S2). Enhancing actomyosin contractility with Rho activator II (1 μg/ml) or nocodazole (10 μM) increased both migration speed and free area fraction by 20-35% relative to untreated controls. Conversely, inhibiting contractility with Y-27632 (20 μM) or blebbistatin (20 μM) reduced both measures by comparable magnitudes. Disrupting cell-cell adhesion with HGF (20 ng/ml) or EGTA (200 µm) also impaired reorganization, with genetic KD-CDH1 producing the strongest reduction (Fig. S5c-d), showing the need for cell-cell mechanical coupling.

### A peripheral mechanical belt of co-enriched E-cadherin and F-actin forms at the colony edge

Parental colonies developed a peripheral belt, a zone of elevated fluorescence intensity extending 25 to 40 μm inward from the lateral colony edges in Zone III (Fig. 2). To determine whether this belt reflects active mechanochemical assembly or passive accumulation of junction material at the tissue boundary, we mapped the spatial distribution of multiple junction components.

**Figure 2.**
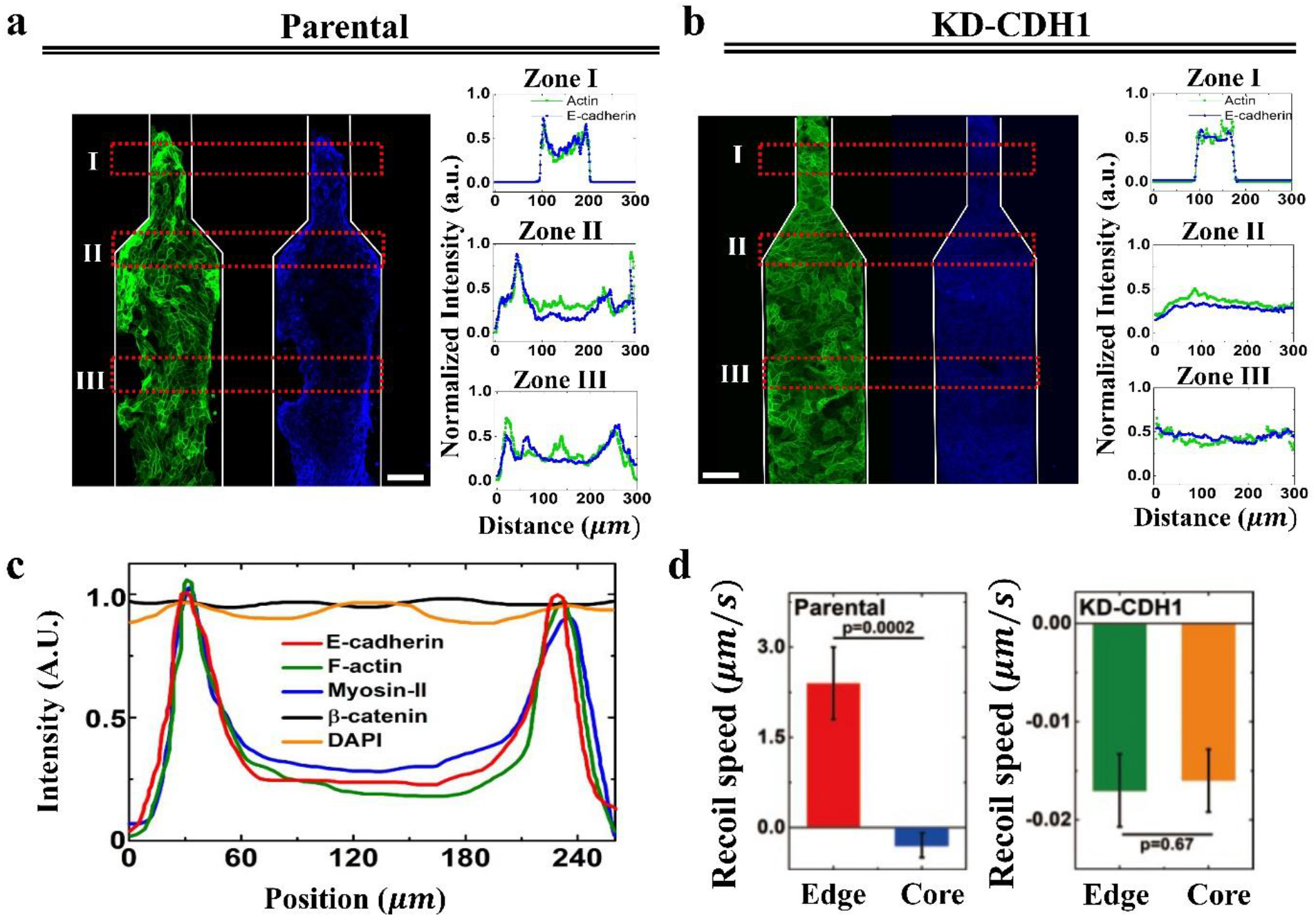
A peripheral belt co-enriched in E-cadherin, F-actin, and myosin-II, but not β-catenin, is under elevated tension. (a,b) E-cadherin (blue) and F-actin (green) in Zone I, II and III of parental (a) and KD-CDH1 cells (b) colonies. Peripheral enrichment of both is present in parental colonies and absent in KD-CDH1. (c) Intensity profiles of E-cadherin, F-actin, myosin-IIA, β-catenin, and DAPI as a function of distance from the boundary of parental colony. (d) Recoil speed after ablation at the edge and core for parental and KD-CDH1 colonies. For statistic evaluation, Welch’s t-test was performed. Mean ± SD, N = 3 to 5 independent batch experiments.

The relative abundance of F-actin and E-cadherin at Zone II belt compared to the center region increased 2.1-fold and 1.9-fold, respectively (Fig. 2a), confirming that enrichment is specific to cell-cell junctions (Fig. S9a-b). In contrast, KD-CDH1 cells showed homogeneous distribution of both E-cadherin and F-actin across the channel width (edge-to-center ratios of 1.1 and 1.05, respectively), with no peripheral belt (Fig. 2b).

Intensity profiles scanned perpendicular to the migration direction showed that myosin-II peaked near the colony edge, with edge-to-center intensity ratios of 1.7 (Fig. 2c; Fig. S9c). β-catenin, an established E-cadherin binding partner, showed no peripheral enrichment (edge-to-center ratio 1.05), remaining homogeneously distributed across Zone III region width (Fig. 2c, Fig. S9c). The absence of β-catenin co-enrichment is inconsistent with a model in which peripheral enrichment reflects passive accumulation of intact junction complexes, which would enrich all cadherin-catenin components proportionally. The data establishes that peripheral and interior junctions differ in their relative composition, with E-cadherin enriched at the periphery independently of β-catenin. Possible interpretations are considered in the Discussion. A plausible self-reinforcing mechanism involves strain stiffening of the actin cytoskeleton under mechanical load (27, 28): peripheral cells under the highest tension would preferentially polymerize F-actin, which in turn stabilizes E-cadherin complexes, further reinforcing the belt. This positive feedback model predicts the sharp spatial boundary we observe but was not directly tested.

Laser ablation directly demonstrates elevated mechanical tension at the peripheral belt (Fig. 2d). Cutting cell stress fibers near the colony boundaries indicated edges caused rapid recoil compared to the interior stress fibers in the core region (Fig. 2d, Fig. S9d). Edge stress fibers in parental cells showed recoil speeds of 2.3 ± 0.7 μm/s, compared to -0.4 ± 0.3 μm/s for core interior stress fibers (*p* < 0.0002, Fig. 2d). This is proportional to local tension when one assumes that viscous resistance is similar across the tissue (29). Stress fiber in the core repaired rapidly without sustained recoil, indicating weak contractile force. KD-CDH1 cells showed uniformly negative recoil rate at both edge (-0.017 ± 0.0037 μm/s) and interior (-0.016 ± 0.0032 μm/s) positions, with no significant edge-core differential (*p* = 0.67; Fig. 2d), consistent with the absence of a peripheral high-tension zone.

Belt formation coincided with extended spatial correlation lengths of cell elongation. Quantified as elongation factor (EF) and F-actin intensity at edge of colony, this is significantly higher than that in the core (Fig. S10a) and the difference becomes even stronger at 20h (Fig. S10b). By 20 h, correlation lengths reached 62 μm for EF and 52 μm for F-actin intensity, corresponding to 5 to 7 cell diameters (Fig. S11; SI Section III). Correlation length increased first in Zone II regions (measurable increase from passage time at 4 h) and later in Zone III regions (measurable increase from 10 h; Fig. S11). This temporal ordering is consistent with retrograde propagation of a coordinated mechanical state from the constriction toward the upstream colony.

### Oscillatory flow dynamically redistributes cell density and prevents bottleneck jamming

PIV analysis, beginning at first contact by leading cells with the narrow inlet, revealed that parental colonies exhibited alternating flow in Zones II and III, those in Zone II advancing and retracting aperiodically (Fig. 1d, Movie S1).

To compare flow and cytoskeletal dynamics on a common scale, we calculated the z-scores of the projected velocity and the F-actin intensity and cross-correlated them within each zone (Fig. 3a-b). For parental cells in Zone II, velocity change precedes the F-actin changes whereas the KD-CDH1 cells require actin intensity change first, then the velocity changes follow, but in Zone I the signals were closely coupled without resolvable time lag, in both lines (Fig. S12).

**Figure 3.**
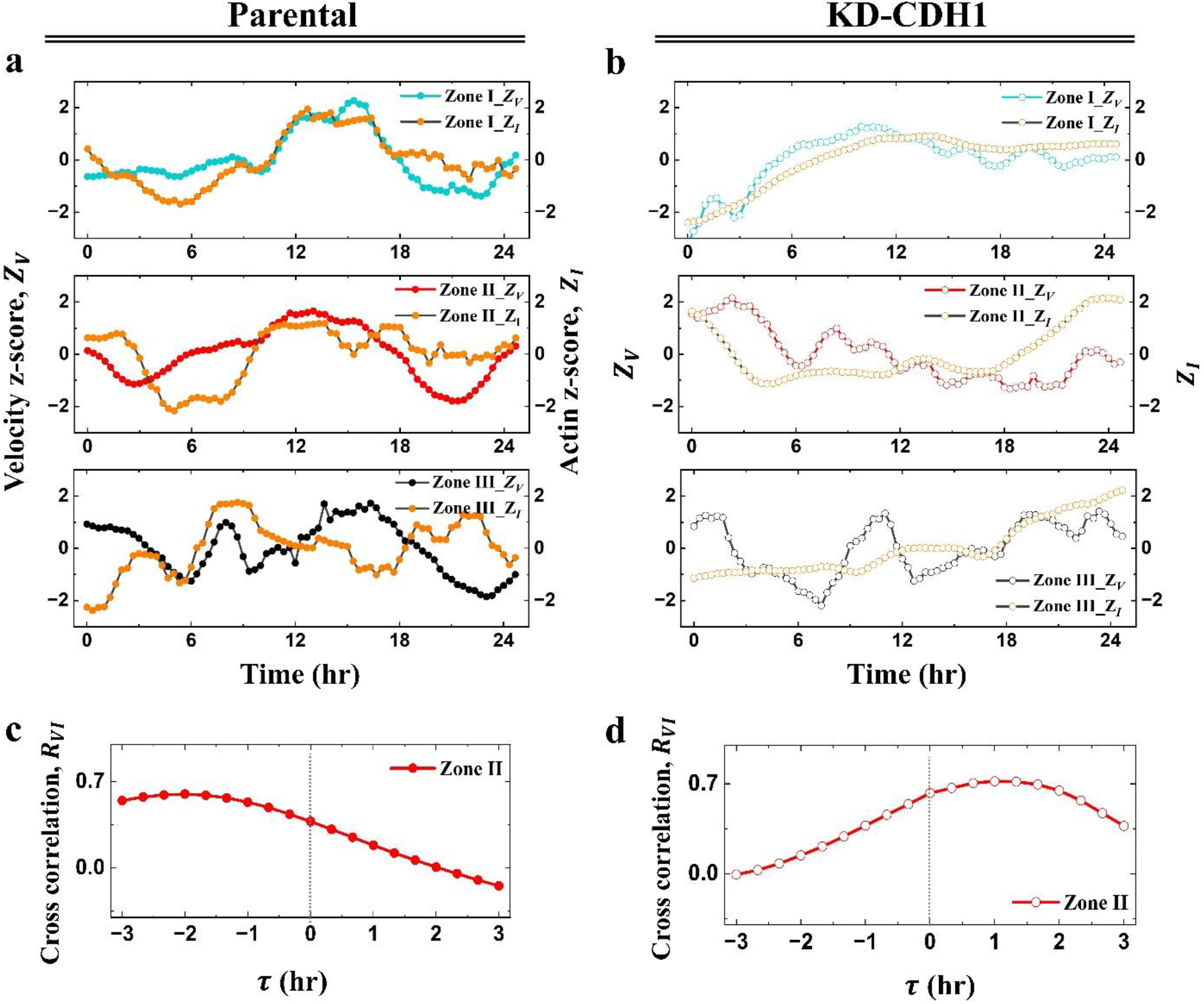
Velocity fluctuations lead F-actin fluctuations in parental colonies but follow them in KD-CDH1 colonies. (a, b) Z-score projected velocity *Z*_*V*_ (filled symbols) and F-actin intensity *Z*_*I*_ (yellow, open symbols), plotted against time in Zones I, II and III for parental (a) and KD-CDH1 (b) colonies as indicated. Time is measured from when the colony enters zone I. (c,d) Cross-correlation *R*_*VI*_(*τ*), of the two signals in Zone II, compared for parental and KD-CDH1 colonies. A peak at *τ* < 0 indicates that velocity fluctuations precede actin fluctuations; a peak at *τ* > 0 indicates the reverse.

We then asked how flow in adjacent zones is coupled, by cross-correlation of projected velocity between Zones I and II, and between Zones II and III (Table S4). Zones II and III correlated more positively in both lines (parental: 0.67 ± 0.13, KD-CDH1: 0.32 ± 0.44) than for Zones I and II (parental: 0.02 ± 0.30, KD-CDH1: 0.27 ± 0.25), indicating that the parental Zone II moves interdependently of the leading front while maintaining coupled to the following cells behind it.

Similar results were confirmed for a sharper geometrical constraint into the narrow inlet, *θ* = 90°, where the switch between forward and retrograde flow in Zone II was more abrupt (Fig. S13).

Oscillational flow was accompanied by density redistribution that relieved bottleneck crowding (Fig. 4). In parental colonies, Zone II initially accumulated the highest density (1750 ± 80 cells/mm^2^), exceeding Zone III density (1450 ± 60 cells/mm^2^) by approximately 300 cells/mm^2^. This we denote as Δφ_*II*−*III*_ (N = 3; Fig. 4a; Fig. S14a). After 10 h, retrograde flow reversed this gradient: cell density in Zone II fell to 1480 ± 70 cells/mm^2^ while in Zone III it rose to 2050 ± 90 cells/mm^2^, yielding a density difference of approximately -600 cells/mm^2^ (Fig. S14a). Simultaneously, Zone I density increased progressively to 1980 ± 110 cells/mm^2^ by 20 h, indicating that cells flowed from Zone II both forward into Zone I and backward into Zone III. Migration speed of parental cells showed large fluctuations across all regions (variation 35-45%; Fig. 4b).

**Figure 4.**
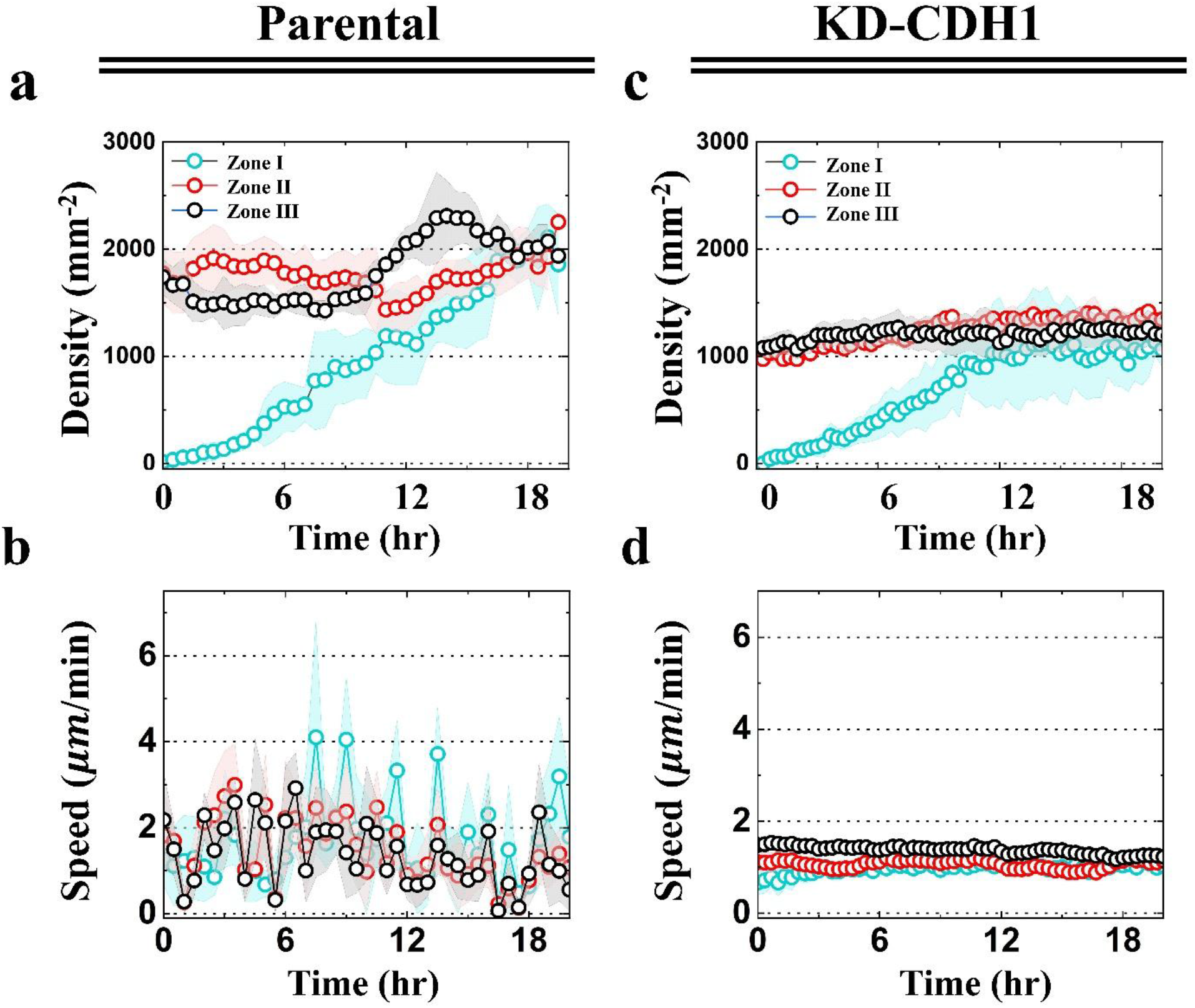
E-cadherin-dependent redistribution of density at the bottleneck. (a,c) Cell density (cells/mm^2^) over time in Zone I (cyan), Zone II (red), and Zone III (black) regions for parental (a) and KD-CDH1 (c) colonies. (b, d) Migration speed as a function of the time for parental (b) and KD-CDH1 (d) colonies, with each region shown in the colors of (a). Shading is SD. Data points are means from N=3 independent batch experiments for each. Time zero is when the colony enters Zone I.

Knockdown of E-cadherin did not show this. We refer to these colonies as KD-CDH1. Overall density was lower (1180 ± 90 cells/mm^2^), differences between Zones remained small, static (Δφ_*II*−*III*_ ≈ ±150 cells/mm^2^; Fig. 4c; Fig. S14a), migration speed was constant and region-independent (Fig. 4d). These static, monotonically graded density profiles are characteristic of passive funneling in which cells compress at the inlet without active redistribution. Oscillatory flow did not reduce transit: the cumulative number of nuclei crossing from Zone II into Zone I over 24 h was 1.5-fold larger in parental than in KD-CDH1 colonies (Fig. S15). This comparison should be read against the lower baseline density of KD-CDH1 colonies (Fig. 4a, 4c).

Proliferation would not explain this, as Zone II density falls over the observation period, and since division can only add cells locally, decrease requires net efflux from that region. This is consistent with the positive flow divergence measured in the same region (Fig. S8b).

### Colony narrowing is preceded by E-cadherin-dependent boundary detachment

Narrowing requires the colony to overcome adhesion to the channel wall, we quantified the competition between peripheral contractility and attachment to the wall, tracking the perpendicular displacement of the colony boundary from the wall, Δ*y*(*x, t*) (Fig. 5a). Boundary fluctuations preceded whole-colony narrowing by 6 to 18 h: local undulations with amplitude 5 to 50 μm appeared at the colony-wall interface before net area reduction was measurable (Fig. 5b-d, Movie S5). In parental colonies these fluctuations grew over time, increasing dramatically after 12 hr, and reaching Δ*y* of 5-120 μm, and eventually gave sustained detachment. In contrast, KD-CDH1 cells maintained stable contact with the channel wall throughout, with boundary fluctuation amplitude below 3 μm (Fig. 5e-g, Movie S5). Nuclei fluctuations in the z direction were also larger at the parental colony boundary than in its interior (Fig. S7c), consistent with local detachment, though it remained below 2µm.

**Figure 5.**
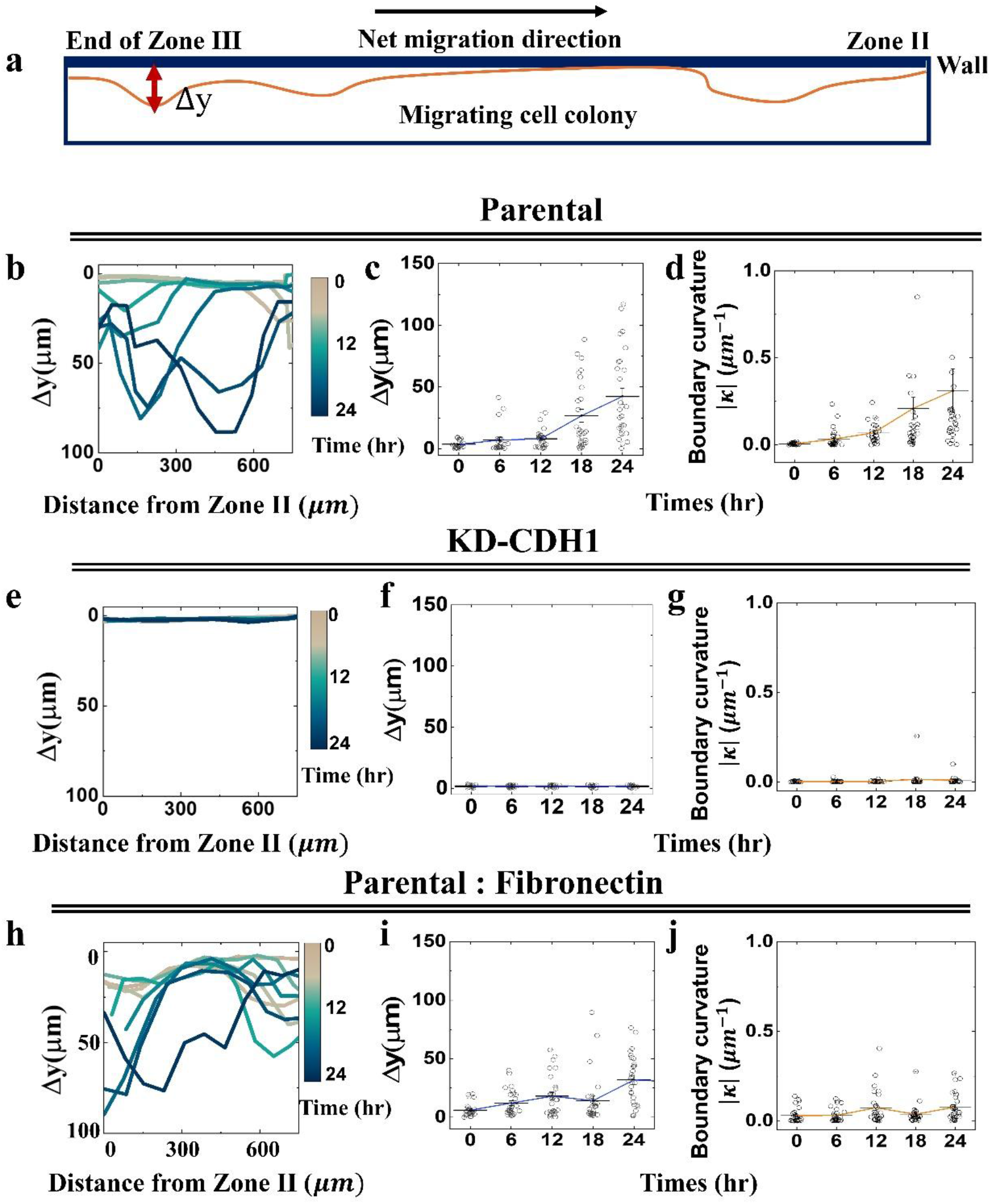
E-cadherin-dependent boundary detachment precedes colony narrowing. Three conditions are compared: parental on uncoated PDMS (b through d), KD-CDH1 on uncoated PDMS (e through g), and parental on fibronectin-coated PDMS (h through j). Time zero is first entry into Zone II. (a) Schematic defining boundary deviation Δ*y*, the perpendicular distance from the colony edge to the straight PDMS wall, as a function of position *x* along the wall. (b, e, h) Deviations Δ*y* at the times indicated by the color scale. (c, f, i) Deviations Δ*y* as a function of time. (d, g, j) Tissue boundary curvature (*κ*) as a function of time; larger *κ* indicates sharper local deformation. N = 3 independent batch experiments per condition.

Detachment was accompanied by sharpening the boundary. We extracted the colony boundary, fitted it with a Bézier curve, and computed the local parametric curvature, |*κ*| (see SI Materials and Methods). In parental colonies, |*κ*| increased from 0.08 ± 0.02 μm^−1^ at 0 h to 0.25 ± 0.06 μm^−1^ at 18 h (Fig. 5d). The largest increase began around 12 h; this coincides with when cell density starts to rise in Zone II (Fig. 4a). The same trend is evident in the time-dependent fluctuation profile, Δ*y*(*x, t*) (Fig. 5c). Higher curvature indicates sharper local deformation as expected for localized contractile force generation. In contrast, the boundaries of KD-CDH1 colonies remained flat with |*κ*| < 0.1 μm^−1^ (Fig. 5g). On walls coated with fibronectin, the onset of wall detachment was delayed, Δ*y* remained within 0 to 100μm, and |*κ*| no longer increased with time (Fig. 5h-j, Movie S5).

These observations together support a working model in which geometric constriction triggers retrograde mechanical signaling through E-cadherin-mediated junctions, induces formation of a contractile peripheral belt, oscillational bidirectional flow that redistributes density, and boundary detachment and colony narrowing.

## Discussion

Powders, colloids and crowds clog when driven through a constriction because their constituents cannot adapt except by contact and do not experience the constriction before reaching it. Adding cohesion makes matters worse, since attraction between constituents promotes arching. Adapting in advance instead requires a collective to sense the constraint, transmit this information across the constituents, and convert it into coordinated behavior change. In this system comprised of migrating epithelial cells, the constraint is the geometric bottleneck and mechanical coupling through E-cadherin junctions carries the information upstream and drives colony-scale adaptation.

We found that colonies develop a peripheral belt as they reorganize under elevated mechanical tension and that belt assembly precedes colony narrowing. Observation of elevated tension at the belt, combined with failure to narrow for E-cadherin knockdown cells that lack the belt, favors a model in which peripheral tension is causally required. While this establishes necessity, the test of sufficiency would require inducing belt formation independent of geometric context, hypothetically by optogenetic activation of RhoA at the colony boundary and observing whether shrinkage of the migrating colony follows. Moreover, the E-cadherin-dependent KD-CDH1 cells exhibit lower steady-state density (approximately 1200 versus 1750 cells/mm^2^ in parental colonies). This density difference may contribute to quantitative differences in migration speed between the cell lines but are unlikely to account for the qualitative absence of oscillational flow, belt formation, and upstream reorganization in KD-CDH1 colonies, none of which shows graded dependence on density in our data.

A central question is whether retrograde reorganization reflects active feedback or passive stress relaxation through a mechanically continuous sheet. In the former, a cell responds to stress by activating a mechanosensitive pathway that generates new stress in its neighbor, which then activates the neighboring cell, so the influence advances upstream at a rate set by cytoskeletal remodeling. In passive equilibration, all cells experience a single stress gradient emanating from the constriction, and the greater delay at greater distances reflects only the finite viscoelastic relaxation of the tissue.

The decisive observation is repeated reversal of flow direction. Inertia is negligible at these scales, so a linear viscoelastic tissue held under a sustained geometric constraint relaxes monotonically towards a steady state and cannot reverse, and the repeated reversals of flow direction we observe are incompatible with that. Timescales suggest the same conclusion. Reorganization advances upstream over 150 to 200 µm in 12 to 17 h, giving 9-17 µm/h for the shape change, which is far slower than elastic stress transmission. MDCK monolayers flowing around a circular obstacle in a confined channel behave as a Maxwell viscoelastic liquid with relaxation time around 70 min (30) in a geometry and cell type close to ours, so passive equilibration should finish within a few hours, whereas our response persists over 6 h. The maximum aperiodic fluctuations at 50 ± 10 µm/h are faster by a factor of roughly four. These distinct observables suggest a fast wave riding on a slowly advancing wave.

The peripheral belt differs from the core in two respects. Mechanically, laser ablation shows it to be under elevated tension; molecularly, E-cadherin is enriched at the belt while β-catenin is not, so the composition of peripheral junctional complexes differs. This raises the question of how the belt transmits force. Measuring α-catenin at the belt, compared with E-cadherin and stress fiber recruitment, could identify the machinery of extended adhesion complexes (31). Another possibility is that the sharp spatial boundary of the belt may reflect tension-dependent stabilization of E-cadherin complexes by F-actin related with supra-cellular actin belt formation (23). These tests, which reach beyond the scope of this study, were not performed.

A specific version of passive equilibration worth inspecting is negative pressure generated by leading cells entering the narrow channel, which would mechanically pull the upstream colony inward. This model makes testable predictions: narrowing should be instantaneous upon leading cells entry, should scale with narrow channel geometry, and should be prevented by increasing cell-substrate adhesion. However, narrowing develops gradually over hours, is independent of taper angle (Fig. S2), and fibronectin coating delays but does not prevent it. Therefore, this explanation does not match our observations.

Cell proliferation during the 24 h observation period could in principle contribute to density changes, but three observations argue against it as the primary driver: the spatial pattern requires net cell transfer between regions, which proliferation cannot produce; the few hours of oscillatory period, which is shorter than the MDCK cell cycle (approximately 18-24 h); and the fact that KD-CDH1 colonies proliferate at comparable rates yet show no redistribution. However, retrograde coupling operates in heterogeneous cell populations, three-dimensional extracellular matrix environments and stiffness, or irregular boundary geometries remain untested, and it is obvious that extension to developmental or cancer contexts would require validation in appropriate cell types and tissue architectures.

The molecular pathway linking mechanical stress at the constriction to cytoskeletal reorganization in upstream cells has not been identified. Candidates include force-induced α-catenin conformational changes, YAP/TAZ-mediated transcriptional responses, and calcium entry through stretch-activated Piezo channels (24). These are not mutually exclusive and likely operate on different timescales: calcium signaling within seconds, conformational changes within minutes, and cytoskeletal reorganization take over hours. That interval is far longer than mechanotransduction at the junction and comparable to the timescale of cytoskeletal remodeling, placing remodeling rather than force sensing as the rate-limiting step. However, mechanosensation and mechanotransduction may determine the magnitude, spatial extent, and temporal persistence of the response, thereby priming the subsequent cytoskeletal remodeling. Beyond direct molecular characterization, epithelial bioelectric activity may provide a complementary means of probing this relationship. Bioelectric features—including spike amplitude, firing frequency, and network-wide propagation—could reveal how local mechanical inputs are converted into spatially coordinated and persistent tissue-scale responses. This possibility is supported by our recent findings that epithelial bioelectric signals are regulated by mechanosensitive ion channels and actomyosin contractility (32, 33).

We conclude that collective cell migration through a constriction is simultaneously a forward and a backward process. Leading cells mechanically signal upstream, and the colony body reorganizes before encountering the geometric constraint. This coupling is carried by E-cadherin-dependent transmission of actomyosin-generated tension and is manifested as a transient peripheral mechanical belt, enabling epithelial colonies to adapt their shape and density distribution to bottlenecks. Following cells are therefore participants in navigation rather than passive recipients of instructions from the front. Cohesion changes how a many-constituent system can respond to a bottleneck.

## Materials and Methods

MDCK epithelial cell colonies were cultured and allowed to migrate through microfabricated confinement channels containing a narrow transition zone; parental and E-cadherin-knockdown (KD-CDH1) cells were analyzed under their respective experimental conditions. Confocal fluorescence imaging was used to quantify collective migration, cellular morphology, actin, E-cadherin and other marker expression, and three-dimensional nuclear distributions. Tissue velocity fields were obtained by particle image velocimetry, and the resulting data were used to calculate axial velocity, flow divergence, zone to zone cross-correlations, and spatiotemporal migration patterns in Zone I (leading cells), II (inlet transition), and III (following cells) regions. Spatial organization of cellular elongation and actin intensity was evaluated using spatial-correlation analyses and corresponding correlation lengths, while colony boundary curvature and cell density quantification and nuclear-position measurements were obtained from segmented image sequences.

Details of experiments for cell culture, channel fabrication, imaging conditions, image processing, quantitative analysis and software are provided in the SI Appendix, Materials and Methods.

## Acknowledgements

Data analysis and writing were supported by startup funds from the Univ. of Massachusetts. We thank Dr. Bo Li for his contribution to live-cell imaging and codes for data analysis at the initial stage of the study.

## Author contributions

S-M.Y., Y-K.C., and S.G. conceived the research. S-M.Y. designed the research, performed experiments, curated, and analyzed the data. S-M.Y. and S.G. wrote the paper.

## Competing interests

The authors declare no competing interests.

**Supplementary information** is available for this paper

## Data availability

The data and analysis code that support the findings of this study are deposited at: https://www.dropbox.com/scl/fo/43caaqt2vs9i606q0ectz/h?dl=0&rlkey=dc7m2pnubjgp2dccmumh0y1v0

