## Supplementary Information for "Mechanical coupling narrows an epithelial colony before it reaches a bottleneck"

### Supporting Information for
### Mechanical coupling narrows an epithelial colony before it reaches a bottleneck

**Sun-Min Yu[a], Yoon-Kyoung Cho[b, 1], and Steve Granick[a, 1]**

[a] Department of Polymer Science and Engineering, University of Massachusetts, Amherst, MA 01003 USA
[b] Department of Biomedical Engineering, Ulsan National Institute of Science and Technology (UNIST), Ulsan 44919, South Korea

[1] **Yoon-Kyoung Cho**


[1] **Steve Granick**


**This PDF file includes:**

Materials & Methods
Supplementary text
Figure S1 to S15
Tables S1 to S4
Legends for Movies S1 to S5
SI References

**Other supporting materials for this manuscript include the following:**

Movies S1 to S5

#### Materials & Methods

##### Channel fabrication

Under conditions designed to produce wall stiffness 2.6 MPa, which is known to promote cell migration, the polydimethylsiloxane (PDMS) microfluidic channels were fabricated by hard and soft photolithography (**Fig. S1**). To fabricate the channel molds, a silicon wafer was spin-coated with negative photoresist (SU-8 2005, MicroChem Inc.), soft-baked, and then exposed under a mask using a contact aligner (MA6, SÜSS MicroTec). Then the hard bake process was followed, and the wafer was soaked in SU-8 developer (KAYAKU advanced materials). The wafer was rinsed with isopropyl alcohol (IPA) to remove residues. To obtain the final channels, PDMS mixture, precursor, and curing agent with mass ratio 10:1 (Sylgard 184 Silicone Elastomer, Dow Corning) was poured onto the wafer and cured at 95 ℃ for 8 h. The peeled-off PDMS channels were directly bonded to the glass substrate of confocal dishes (SPL Life Science 101350) after plasma treatment (Cute-MP, Femto Science) under 50 sccm oxygen and 80 W for 90 s.

For surface coating with fibronectin, confocal dishes having PDMS microfluidic channels were fully filled with 10 μg/ml of fibronectin (F0895; Merck) in PBS for 2 h at 25 ℃.

##### Cell line maintenance

The MDCK-WT cell line (Madin-Darby canine kidney wild type, MDCK NBL2; ATCC) was cultured and seeded. Cells were cultured in low-glucose DMEM (11885084; Thermo Fisher Scientific) with 10 % fetal bovine serum (16000044; Thermo Fisher Scientific) and 1 % penicillin-streptomycin (15240062; Thermo Fisher Scientific), in a humidified 5 % carbon dioxide incubator at 37 ℃. For cell culture maintenance, media was changed every two days, and cells were sub-cultured at nearly 90 % confluence using 0.25 % trypsin (25200072; Thermo Fisher Scientific). Harvested cells were dispended in a confocal dish (101350; SPL) having PDMS microfluidic channels with a seeding density of 3.2 × $10^4$ $cm^{-2}$. Three days after seeding, the cells reach a confluent state outside the wide channel and start to migrate. At this time, we put the sample on the microscope stage and start taking videos. For MDCK-KD-CDH1 cells, culture medium containing 2 μg/mL tetracycline was introduced 48 h before actual live-cell imaging began.

##### Cell line development for live-cell imaging

The plasmids are listed in **Table S1**. To visualize F-actin and nucleus (Histone H2B), the following stable fluorescent cell lines were used: MDCK-H2B-mCherry and Lifeact-mEGFP, obtained as plasmid gifts from Robert Benezra and Michael Davidson, respectively (#20972 & #54610; Addgene). Co-transfection enabled us to simultaneously monitor the nucleus and actin cytoskeleton, following the evolution of mechanical signal, and shape of the cell body and nucleus, along with dynamics and morphology, all of this *in situ*.

The cell transfection was conducted using the 4D-Nucleofector system (AAF-1002B; Lonza) with SE cell line 4D-Nucleofector X kit (V4XC-1024; Lonza). Afterwards, G418 antibiotic selection (Geneticin, 10131035; Thermo Fisher Scientific) was carried out at 0.6 mg/mL to obtain stable cell lines. Single clones were selected that showed stable expression of both Lifeact-mEGFP and H2B-mCherry.

##### CDH1 knockdown cell model

To assess the influence of E-cadherin adherence junction protein, we developed a monoclone having knockdown expression of the CDH1 gene. To accomplish this, shRNA expression against CDH1 gene was designed to be operative to tetracycline induction by cloning oligonucleotides of shRNA-CDH1 into the AgeI/EcoRI site of pLKO-TetOn-puro (#21915, Addgene) as follows;

Forward:5'–CCGGGCTCTCATTTCCGATTATATTCTCGAGTTCGAGAGTAAAGGCTAATATTTTTG–3'

Reverse: 5'–AATTCAAAAATATTAGCCTTTACTCTCGAACTCGAGAATATAATCGGAAATGAGAGC–3'.

The titration of lentivirus particle was measured by p24 ELISA test and original stock concentration was measured as 2.61x$10^8$ TU/ml. The virus particles were transduced to the parental cell line (MDCK-H2B-mCherry & Lifeact-mEGFP) in the presence of 5 μg/ml polybrene. The infected cells were selected for 3 weeks in culture media containing 3 μg/mL puromycin. The final monoclone cells MDCK KD-CDH1 cells (MDCK-H2B-mCherry & Lifeact-mEGFP & pLKO-TetOn CDH1-shRNA) were incubated with 2 μg/mL of tetracycline for 48 h to have sufficient knockdown (T7660; Sigma-Aldrich). The level of E-cadherin was confirmed by Western blot analysis of protein extracts.

**Western blots**
Proteins for MDCK cells were extracted using RIPA buffer with 1X cOmplete Protease Inhibitor cocktail (11697498001; Merck). The supernatants of protein lysates were centrifuged at 16,000 g at 4 ℃ for 20 min. Protein concentrations were quantified by BCA protein assay (23250; Thermo Fisher Scientific). For denaturation, the protein solution was mixed with the sample buffer solution and reducing agents, then the mixture was heated at 95 ℃ for 10 min. The protein (30 μg) was loaded onto NuPage 7% Tris-Acetate gel using a mini gel tank and wet transferred onto polyvinylidene fluoride (PVDF, pore size: 0.45 μm) membrane using a Mini Blot module (NW2000; Thermo Fisher Scientific). The PVDF membrane was blocked by 3% non-fat dry milk in TBST buffer (Tris-buffered saline containing Tween 20). The primary antibodies were incubated against E-cadherin (1:1,000 dilution, rat anti-E-Cadherin antibody; DECMA-1, ab11512; Abcam) and α-tubulin (1:2,000 dilution, mouse anti- α-tubulin antibody, T5168, Merck) at 4℃ overnight incubation. The blots were then washed 3-4 times for 10 min each in TBST. They were then incubated with Alexa-647-linked secondary antibodies against rat-IgG (1:10,000 dilution, ab150159, Abcam) and mouse-IgG (1:10,000 dilution, A32728, Thermo Fisher Scientific). The blots were then washed three times with TBST for 10 min each. The blots were then revealed using CHEMIDOC MP (BioRad).

**Pharmacological interventions**
The pharmacological chemicals are listed in **Table S2.** Pharmacological interventions were performed to up- and down-regulate actomyosin contractility. The cells were treated with 20 μM Rho kinase inhibitor Y-27632 (SCM075; Merck) and myosin II inhibitor Blebbistatin (B0560; Merck) for 6 h. To enhance actomyosin contractility, cells were treated with 1 μg/mL Rho activator II (CN03-A; Cytoskeleton) for 3 h. To perturb cell-cell adhesion interaction, cells were treated with 20 ng/ml hepatocyte growth factor (HGF) (H9661; Merck) and 200 μM ethyleneglycol-bis (β-aminoethyl)-N,N,N',N'-tetraacetic Acid (EGTA) (03777; Merck) for 3h. To induce microtubule depolymerization and thus indirectly affect actin polymerization, the cells were treated with 10 μM Nocodazole (M1404; Merck) for 6 h. Afterwards, confocal live-cell imaging was conducted.

**Immunocytochemistry**
To achieve staining of cells inside the channels, all incubation steps were performed on a shaker horizontally rotating at 60 rpm. Cell samples were fixed with 4 % paraformaldehyde for 20 min following gentle PBS washing. Fixed cells were permeabilized with 1 % Triton-X in PBS for 20 min and washed twice with PBS for 10 min each. A blocking step was performed by exposure to 5 % bovine serum albumin (BSA) in PBS solution for 1 hr. Phalloidin-fluorescein isothiocyanate (FITC; P5282; Sigma-Aldrich) was applied for a 40 min incubation at 1:500 dilution. For nucleus staining, cells were stained with 4′, 6-diamidino-2-phenylindole (DAPI) (D9542; Sigma-Aldrich). The primary antibody incubations were performed at 4 ℃ overnight incubation in 1 % BSA in PBS; primary anti-E-cadherin antibody (DECMA-1) (1:100 dilution, ab11512; Abcam), primary rabbit anti-β-catenin antibody (1:200 dilution, ab16051, Abcam), and primary rabbit anti-myosin-II antibody (1:200 dilution, M8064, Merck). Secondary antibody labeling with goat anti-rat IgG antibody-Alexa405 (1:500 dilution, ab175671; Abcam) and goat anti-rabbit IgG-Alexa 647 (1:1,000 dilution, A32733, Abcam) was performed in 1 % BSA in PBS for 2 h at room temperature each.

**Live cell confocal microscope imaging**
The cells were kept in an incubator to maintain an environment of 5 % carbon dioxide and 37 ℃ throughout all imaging processes. For confocal imaging, we used a laser scanning confocal microscope (FV3000, Olympus) equipped with 20x phase contrast objective (UCPLFLN20X). The total imaging time was set to 48 h with a frame rate of 3 or 2 frames per hour, at three different positions capturing 1900 μm × 640 μm for single full microchannel with the x and y pixel resolution

as 0.62 $\mu$m. The wavelengths of the exciting laser were selected as 488 nm (mEGFP) and 561 nm (mCherry) to image F-actin and the cell nucleus, respectively. To image DAPI signal in stained samples, the excitation wavelength was 405 nm. At each position of imaging, six z-scans were performed from the apical to the basal side of the cells with vertical steps of 3 $\mu$m. Each image is reconstructed from a stack of six confocal images scanned with this procedure.

**Laser ablation**

The laser ablation experiments were conducted using a laser microdissection system (PALM Microbeam, Zeiss) equipped with pulsed FT-UV laser. To cut the edge of a cell colony, we focused a ns pulsed 355 nm UV laser on the selected spot for 0.5 s. The small beam size, 0.6 $\mu$m in diameter, ensured that perturbation caused by the laser was localized to the edge junction without influencing the cell nucleus. The 1-2 ns duration of each laser pulse effectively avoided heat accumulation within the cell. Immediately after laser ablation, *in situ* live cell imaging automatically started at the same platform at 0.5 fps. The cells were kept in an incubator to maintain the environment of 5 % carbon dioxide and 37 ℃ throughout the imaging processes. A 40x phase contrast lens (Korr LD Plan-Neofluar) and color CCD (AxioCam Icc1 Rev. 4) were used to capture the fluorescence images.

**Quantification of images**

The discrimination of individual cells and digitization in the plane were performed using IDL code written by us. Since F-actin usually localizes near the plasma membrane, it serves as an indicator of the cell boundary, so we used the F-actin channel of the confocal images for cell segmentation. We first gray-scale each image pixel to the range of 0-255. To ensure that the center of the cells is the bright spot in the image, we revert the image by subtracting 255 from each pixel value. We then apply the watershed algorithm to produce the segmented cell boundary. The algorithm first finds local maxima of the image and pre-features the central region of the cells before many fastest descendant operators are applied simultaneously to each local maximum. This procedure causes the operators to move in the direction in which the brightness, or equivalently the F-actin signal strength, decreases most rapidly. The positions where two or more operators meet are marked as the boundaries of the cells.

The above watershed procedure gives the pixel coordinate information of the cell boundaries. To sort the boundary pixels to each cell, we additionally conducted the following steps. For each cell, starting from the local brightness maximum corresponding to it, we construct a ray from it. The first boundary pixel that meets with this ray is regarded as belonging to the boundary of this cell. We then rotate the ray from $0^\circ$ to $360^\circ$ degrees with steps of $0.2^\circ$and thus find all the boundary pixels for this cell. This procedure is repeated for all cells. A polygon fit is then applied to the boundaries to decide the region that they enclosed, and thus the inner cell regime is digitized.

For 3D image analysis, we used object detection algorithms and tracking modules in Aivia AI Image Analysis Software (version 16.0, Leica Microsystems, Germany) to track fluctuations in the z-axis direction. Individual detected nuclei were reconstructed in 3D space based on analysis of their x, y, and z coordinates to deduce volumetric morphological changes. To classify 3D fluctuations of cell nucleus positions in different regions of the migrating cell colonies, we classified cross-sections of the migrating cells according to their distance from the colony central line such that 30% resided in the interior (“core”) and 30% at each of the edges, and rest of section between two regimes was classified as intermediate (**Fig. S7c**). The z-coordinate of each nucleus was adjusted by subtracting the mean z-coordinate of all nuclei detected at $t$ = 0. These values are shown as adjusted nuclear z-coordinates in Fig. S7.

**Morphological information**

Morphological information, such as the area and perimeter of the cell and nucleus, is read from the number of pixels on the boundary or enclosed by the boundary. We apply ellipsoidal fitting to the boundary points and use the aspect ratio and orientation of the fitted ellipses as the non-sphericity and orientation of the cells, respectively. We defined elongation factor (EF) as the ratio of the length of the long axis to the short axis. We also detect the intensity of each marker (F-actin, DAPI, H2B, E-cadherin) within the cell boundary by reading the intensity of individual color channels: green for F-actin, red for H2B, cyan for DAPI, blue for E-cadherin.

As DAPI stains DNA, it reflects the morphology of the nucleus. We therefore select pixels within the boundary whose intensity exceeds a threshold (usually 40 on the 0-255 color scale) and mark them as belonging to the nucleus. We do the same counting and fitting to pixels that belong to the nucleus and obtain morphology information about the nucleus. From this procedure, we extract information of each cell into a fourteen-column list (1): [$\langle x_{cell} \rangle$, $\langle y_{cell} \rangle$, $\theta_{cell}$, cell inverse aspect ratio, F-actin intensity, E-cadherin intensity, $\langle x_{nucleus} \rangle$, $\langle y_{nucleus} \rangle$, $\theta_{nucleus}$, nucleus area, nucleus inverse aspect ratio, DAPI intensity, time, cell ID]. From this data, we analyze statistics of various quantities in the main text and study their time evolution.
The analysis codes for cell morphology analysis are deposited at: https://www.dropbox.com/scl/fo/43caaqt2vs9i606q0ectz/h?dl=0&rlkey=dc7m2pnubjgp2dccmumh0y1v0.

When F-actin is not distinctly localized at the cell boundary, the accuracy of cell segmentation can be unsatisfactory. In these instances, we analyze the image based on tracking the nucleus. We extract the nucleus channel from the confocal image and enhance the contrast using the built-in adjustment function in ImageJ (NIH Java 1.8.0.112). Then we use a connectiveness method (home-built MATLAB script) (2) to identify the position and orientation of the nucleus. The averaged F-actin (or E-cadherin) strength is then the sum of the brightness in all pixels normalized to the number of nuclei identified.

To quantify the spatial distribution of cell density, we coarse-grain each image by dividing it into equal square boxes, each 35 $\mu m$ × 35 $\mu m$. We then count the number of cells within each box. For a cell that lies on the boundary of a box, we calculate the area ratio between those portions inside and outside the box. If the ratio is greater than one, we count this cell as being inside the box; and conversely if the ratio is less than one. Finally, we average the values for all boxes with the same x-position (**Fig. 4, Fig. S14**)

**Curvature measurement of tissue boundary**
The boundary of the migrating tissue fluctuated in Zone 3 with local curvature. Time-lapse images were converted to gray scale, thresholded on the green fluorescence actin signal, and segmented as a single object. The sign of curvature is determined by vector direction from the osculating circle, inside or outside the migrating tissue, but we report the absolute value to emphasize amplitude rather than direction. Curvature was defined as follows, where the first is for a circular arc of constant curvature, and the second is a parametrized boundary of varying curvature (our situation):

$$|\kappa| = \frac{1}{R}$$

or

$$|\kappa| = \frac{|\dot{x}\ddot{y} - \dot{y}\ddot{x}|}{(\dot{x}^2 + \dot{y}^2)^{3/2}}$$

Curvature at each point along the cell boundary was calculated from the latter equation where $x(t)$ and $y(t)$ are the Cartesian coordinates of the boundary contour parameterized by $t$, and $\dot{x} = dx/dt$, $\dot{y} = dy/dt$ denote the first derivatives and $\ddot{x} = d^2x/\mathrm{d}t^2$, $\ddot{y} = d^2y/\mathrm{d}t^2$ denote the second derivatives with respect to $t$. Curvature was extracted using the Kappa plug-in in the FIJI implementation of ImageJ(3), manually defined by control points and continuous curvature extracted at 1,000 points per contour.

**Particle image velocimetry (PIV) analysis**
Velocity fields were obtained using particle image velocimetry, MATLAB PIVlab (version R2026a) (4). Each 1900 μm × 640 μm for single full microchannel with the x and y pixel resolution as 0.62 μm was analyzed in three iterations with interrogation windows of 64 × 64 (40 × 40 $\mu$m), 32 × 32

(20 × 20 $\mu$m), and 16 × 16 pixel (10 × 10 $\mu$m). Vectors from the final 16 × 16-pixel iteration are reported. Frames were acquired every 20 or 30min, giving a velocity resolution of 0.1-0.2 μm/h were then visualized by arrow plots reflecting both the direction and magnitude of the velocity.

To represent motion along the migration axis, we averaged the x-component of the velocity field over y at each x and each time point, giving the axial velocity, $V(x,t)$. Positive values indicate forward motion, negative values the reverse. Time zero is first leading cells contact with the inlet. Spatial position was measured as defined in Fig. 1a.

**Divergence of the velocity field**

The two-dimensional divergence, reported here in $h^{-1}$, identifies local regions of net outflow ($\nabla \cdot \mathrm{v} > 0$) or the reverse. With $u$ and $v$ the x and y components of the PIV velocity field,

$$\nabla \cdot \mathrm{v(x, y, t)} = \frac{\partial u(x, y, t)}{\partial x} + \frac{\partial v(x, y, t)}{\partial y}$$

Divergence was averaged over each region,

$$\langle \nabla \cdot \mathrm{v} \rangle_{\Omega}(t) = \frac{1}{A_{\Omega}} \int_{\Omega} \left[ \frac{\partial u(x, y, t)}{\partial x} + \frac{\partial v(x, y, t)}{\partial y} \right] dA,$$

Where $\Omega$ denotes the region of interest and $A_{\Omega}$ is its area.

**Standardization and cross-correlation of velocity and F-actin intensity**

To correlate changes in velocity and stress fiber intensity, we performed z-score standardization. The PIV vector magnitudes and directions were denoted by $\mathrm{m}(t)$ and $\theta(t)$, respectively. The projected velocity components were aligned with the net migration axis as

$$V_{\parallel}(t) = \mathrm{m}(t) \cos( \theta(t) - \theta_{axis}),$$

where $\mathrm{m}(t)=\sqrt{u(t)^2 + v(t)^2}$ and $\theta_{\mathrm{axis}}$ the direction of net colony migration.

Because $V_{\parallel}$ and $I$ carry different units, both were standardized before comparison. Standardization gave

$$Z_V(t) = \frac{V_{\parallel}(t) - \overline{V_{\parallel}}}{\sigma_V} \text{ and } Z_I(t) = \frac{I(t) - \bar{I}}{\sigma_I},$$

where $\overline{V_{\parallel}}$ and $\bar{I}$ indicate the mean of the $V_{\parallel}$ and $I$ and $\sigma_V$ , $\sigma_I$ are their standard deviations. The normalized cross-correlation is

$$R_{VI}(\tau) = \frac{\sum_i [Z_V(t_i + \tau)][Z_I(t_i)]}{\sqrt{\sum_i Z_V(t_i)^2 \sum_i Z_I(t_i)^2}}$$

where $\tau$ is lag time. A peak at $\tau < 0$ indicates that velocity fluctuations precede F-actin fluctuations, and the converse.

**Statistical validation**

Comparisons between two independent groups were performed using a two-sided Welch's two-sample t-test in OriginPro 2023b (OriginLab). This test does not assume equal variances between groups. Data are presented as mean ± standard deviation (SD), unless otherwise specified. The number of independent experiments (N) and *p*-values are provided in the corresponding figure legends. Differences were considered statistically significant at $p < 0.05$.

#### Supporting Information Text

##### I. Detailed description of the sample configuration.

Here we offer more information about the sample configuration for cell migration (**Fig. S1**), through channels that were produced by photolithography. The channel length is 2 mm. By using different types of the photoresist and adjusting the spin coating rate, we fabricate channels with thickness from 20 to 150 $\mu\mathrm{m}$. After seeding the cells onto the free areas of the confocal dish, the cells first reach confluence outside the channel area and then migrate into the entrances of the channels from both sides (red arrows in **Fig. S1a**) in response to the attraction of free area inside. This symmetric design with two open entrances ensures free diffusion of nutrients in the culture medium within the whole sample and therefore the same chemical environment inside and outside the channel. This reorganization finishes within 24 h after the cells enter the narrow inlet channel, well before the cells from the two sides meet in the middle of the narrow channel. Therefore, migration from the two entrances are independent events without mutual interference. This excludes the hypothetical possibility that reorganization is caused by a dragging force from the opposing colony when cells from both sides meet. Three independent biological replicates dataset are prepared.

In principle, one may worry that the ceiling of the channel may affect the metabolism of the cells beneath it due to depletion of cell medium or alteration of signaling via soluble cytokines(5, 6). However, our sample configuration is designed to avoid cell medium depletion. The smallest channel heights in our experiments, 20 $\mu\mathrm{m}$, much exceeds the apical-basal height of the MDCK cells (less than 10 $\mu\mathrm{m}$) leaving sufficient z-space above the migrating colony. This ensures that medium inside the migration channel can exchange with that outside efficiently.

**Fig. S2** shows the data from which we determined the optimal width of the inner inlet and the inlet angle. This figure also shows that height of the PDMS ceiling does not matter. In control experiments conducted under different channel heights, none of the quantities we measured show significant change. Even considered on the molecular level, the strength of F-actin expression is unaffected. On the cellular level, the cell density does not depend on channel height. In addition, no sign of 3D structure formation was observed in either pristine or fibronectin coated samples. This justifies that the height range (20-150 $\mu\mathrm{m}$) we chose is suitable for the in-plane migration of the epithelial tissue.

One may legitimately wonder about the influence of substrate stiffness. Surveying the literature, we note that glass substrate proves to be a suitable condition for study of collective migration in epithelial systems (7-10). Quantification on flat surfaces shows lesser uniform forces on glass than on substrates with lesser stiffness (10). In contrast, we observe enhanced F-actin expression starting from the tapering corner. Therefore, the migration scenario described here is not caused by substrate stiffness.

##### II. Extended discussion of pharmacological interventions.

Building upon information from the literature (1,11-16), we conducted pharmacological interventions to these cell colonies in the confluent, non-migrating state (**Fig. 2**) using the same MDCK cell line as in our cell migration experiments. This offered the chance to test the migration scenario reported in the main text by switching, on and off, key components one at a time. We summarize the outcome of these pharmacological interventions in **Table S2**. The consequences of pharmacological intervention are consistent with the reorganization scenario proposed in the main text.

At first glance, the outcome of HGF and EGTA may look surprising. HGF is motogenic, in addition to mitogenic, so epithelial cells treated with HGF acquire additional motility. Normally, this facilitates movement-related processes such as migration, morphogenesis and acquisition of invasiveness via upregulation of Snail (EMT marker) (14,15), but in our experiments, HGF is observed to impede migration (**Fig. S5c**), which may seem counterintuitive. The apparent reason is that HGF reduces the degree of confluence or packing density of cells by weakening the cell-to-cell interaction; this is why HGF is also called 'scatter factor'(16). Loss of collective response within the epithelial colony impedes whole-colony reorganization. We conclude that the observed HGF action is consistent with our interpretation of whole-colony collective response.

#### III. Calculation of spatial correlations.

To evaluate the spatial correlation of the morphological parameters of epithelial cells, we calculated the correlation coefficient value from Moran's *I* spatial correlation using the formula: $\mathrm{g}(\psi) = \frac{n}{W}\frac{\sum_i^n \sum_j^n w_{ij}(\psi_i-\bar{\psi})(\psi_j-\bar{\psi})}{\sum_i^n(\psi_i-\bar{\psi})^2}$, where *n* is number of samples, $\psi_i$ and $\psi_j$ indicate the values of the variables (EF and F-actin strength) at locations *i* and *j*, respectively, $\bar{\psi}$ is the mean of variable, $w_{ij}$ is a matrix of weighted values, where elements are a function of distance, and $W$ is the sum of the values of the matrix $w_{ij}$. The space correlation, $\mathrm{g}(\psi)$, is zero for no correlation and 1 for a perfect association between variables.

We calculate spatial correlation in Zones II and III at different times. When cells enter the tapering inlet, $\mathrm{g}(\psi)$ is greatly enhanced for both quantities at Zone II, indicating the emergence of long-range correlation. At later times, $\mathrm{g}(\psi)$ of Zone III cells start to increase. Note that the quantities we measured -- cell elongation, F-actin intensity, and so forth -- all show heightened correlation, which suggests that the increase is a property of the tissue.

#### IV. The alternative hypothesis of cavitation.

As an alternative to the mechanical feedback scenario proposed in the main text, one might argue that as the leading cells speed up, they exert force on cells behind them to produce tension not only along the direction of the channel but also orthogonal to it. By this argument, a cavity opens in the same way that cavitation bubbles form in a liquid whose pressure is sufficiently negative (17).

However, increasing cell adhesion to both the PDMS walls and the glass substrates by treating them with fibronectin produced no qualitative change: the average free area ratio showed no change within error bars defined as standard deviation of the measurements (**Fig. S2a, b**). It appears that the two-dimensional contact between cells and the glass beneath them provides mechanical stability to counterbalance potential forces caused by this argument. Moreover, accumulation of F-actin in Zone II makes these cells more rigid than those upstream, which is inconsistent with the cavitation-based assumption that the migrating colony is a simple fluid. Likewise inconsistent is that the optimum narrow channel width ($w'$) is observed to be independent of the taper angle $\theta$; in a hypothetical negative pressure scenario, its magnitude would scale as the inverse of $\theta$ and the optimum $w'$ would have negative correlation with $\theta$. Contrary to such an argument, we observe the same optimum $w'$ regardless of $\theta = 45^\circ$ and $60^\circ$ (**Fig. S2c**).

A third reason to doubt the cavitation scenario is our observation that significant phase differences emerge from inspecting time traces of different signals, unlike the homogeneous response anticipated for a cavitation process. We observe that the leading cells start to accelerate in response to contact with the tapering corner and that colonies in Zones II and III reorganize.

#### Figures

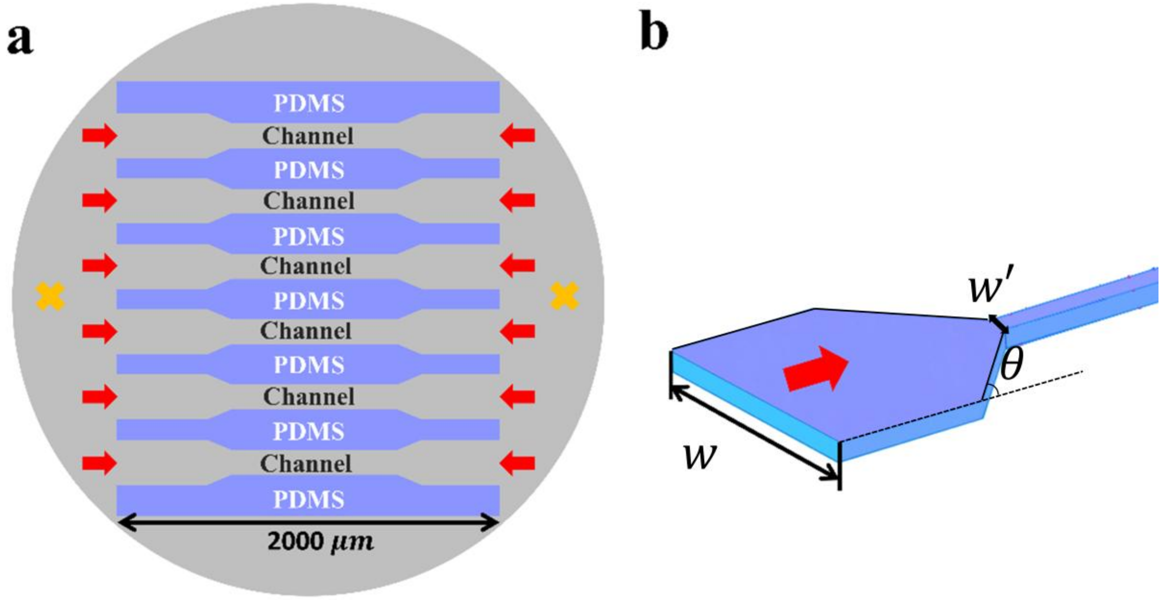


**Figure S1. Microchannel geometry and experimental configuration.** (a) Schematic layout. Grey circles represent the glass area of the confocal dish; orange crosses, cell seeding positions; red arrows, direction of migration. (b) Three-dimensional schematic of the PDMS channel in which cell colonies migrate, with $w$ and $w'$ indicating the width of the wide and narrow straight channel, respectively and $\theta$ the taper angle of the inlet.

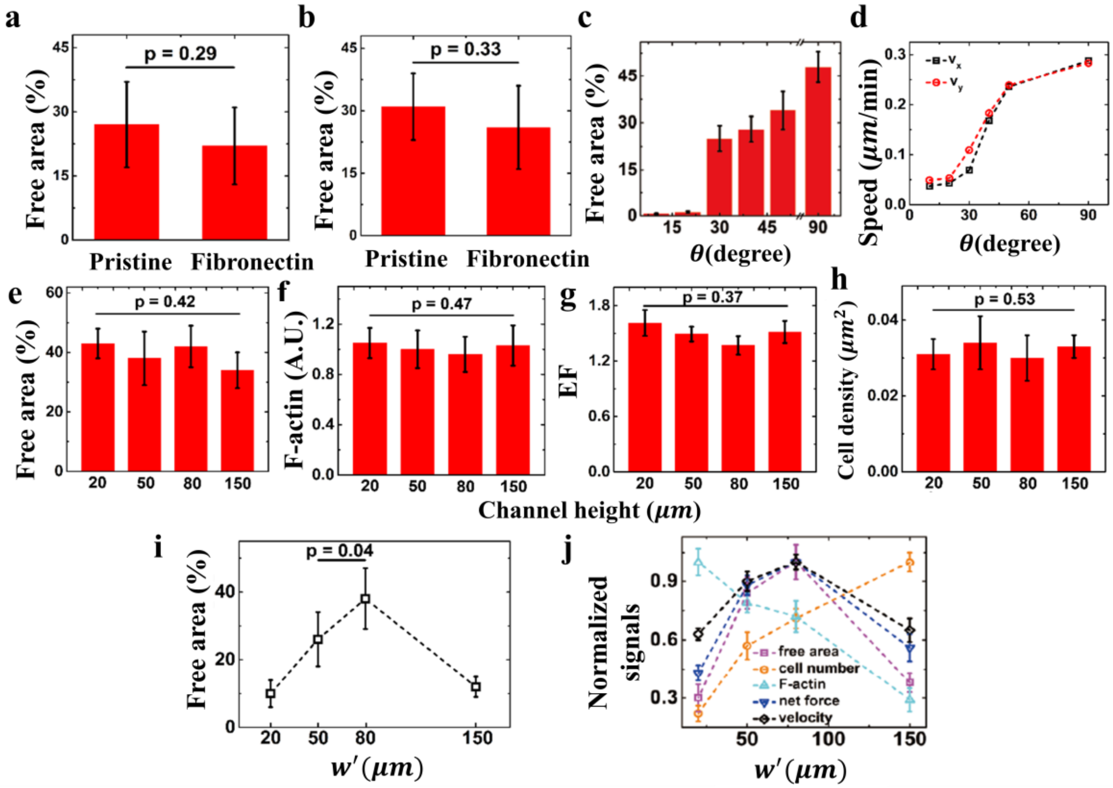


**Figure S2. Determination of optimal channel fabrication parameters.** Student's *t*-test was performed between the pristine and fibronectin-coated situations. The error bars indicate the standard deviation. (a) Comparison of pristine PDMS channel walls and channel walls treated with fibronectin to increase cell-wall adhesion for $\theta$ = 45° (a) and 60° (b). For untreated PDMS and fibronectin-treated PDMS, these bar charts show the averaged area fraction (N = 5) at $t$ = 20 h and inlet width $w'$ = 30 $\mu\mathrm{m}$. (c) For the untreated PDMS channels, the average free area fraction (N = 5) is plotted as a function of inlet taper angle. (d) Plots obtained from particle imaging velocimetry (PIV) show that the average speed of cell movement increases with increasing taper angle in both x and y directions. (e-h) Channel height dependence. None of the measured variables, including free area (e), F-actin strength (f), cell elongation factor (EF) (g), and cell density (h) exhibit significant changes when the cells migrate in channels with heights varying from 20 $\mu\mathrm{m}$ to 150 $\mu\mathrm{m}$.(i) The inlet taper width ($w'$) dependence of free area fraction at 20 h and $\theta = 45^{\circ}$. (j) The width ($w'$) dependence of various signals indicated in the figure panel, at 20 h and $\theta = 45^{\circ}$. Error bars are the standard deviation from 10-20 samples in i and j.

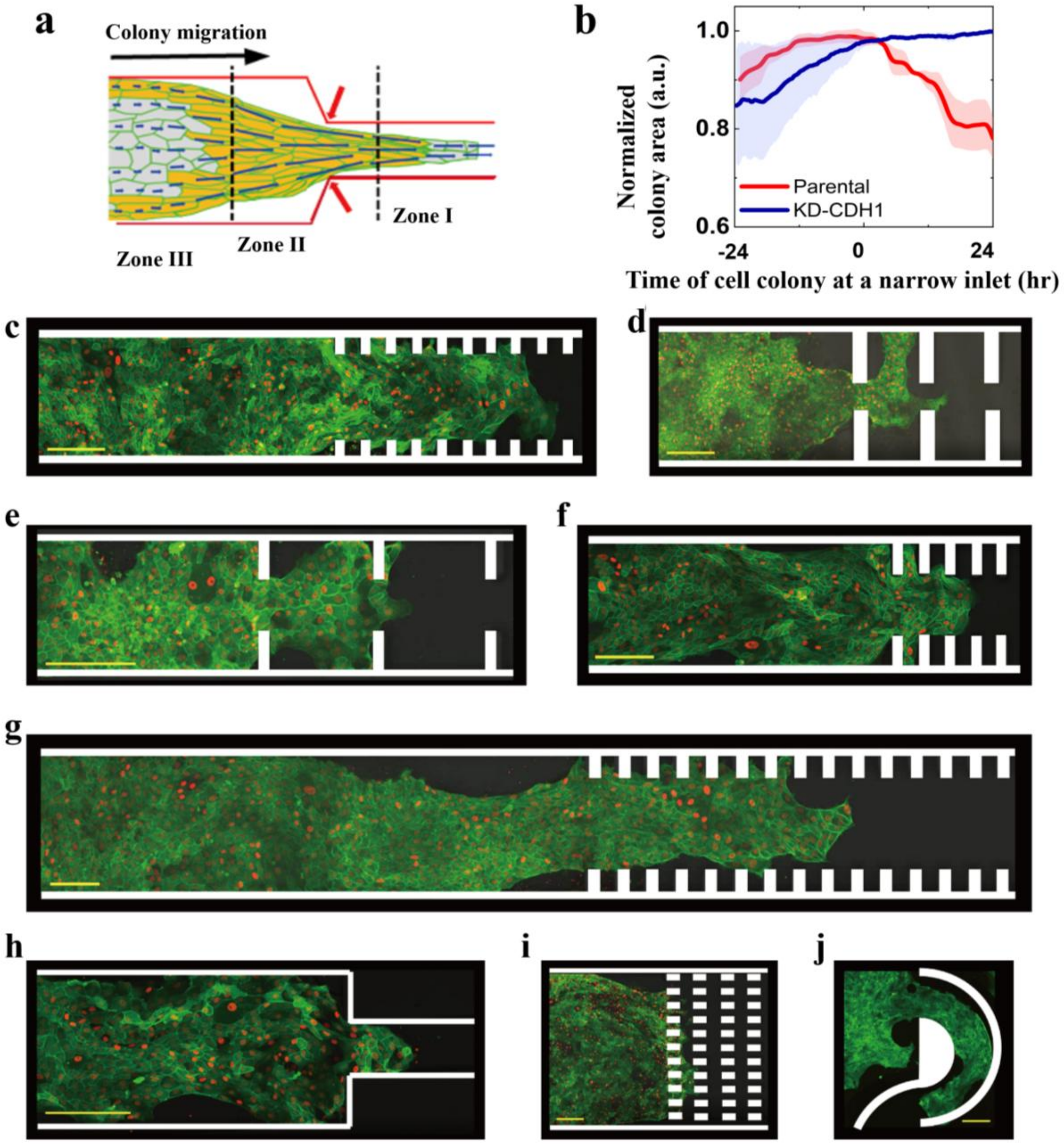


**Figure S3. Generality of wall detachment for additional geometries.** (a) Standard geometry. A colony migrates from a wide channel ($w$ = 250 μm) through a tapered inlet ($\theta$ = 30°) into a narrow channel ($w'$ = 80 μm). Zones I, II and III are defined by spatial position relative to the inlet. Yellow shading: peripheral belt. (b) Normalized colony area over time for parental and KD-CDH1 colonies in the microchannel with standard geometry. N=4; shading: SD. N is the number of independent experiments. (c-j) Fluorescent confocal images of the MDCK-H2B-mCherry & Lifeact-mEGFP cells colonies migrating in channels with protruded bar array along the straight channel wall (c-g), 90-degree tapering (h), rectangular array (i), and curvilinear (j) geometry. Green denotes F-actin expression, red denotes the cell nucleus. Channel shapes are indicated in white. Scale bar, 100 $\mu m$.

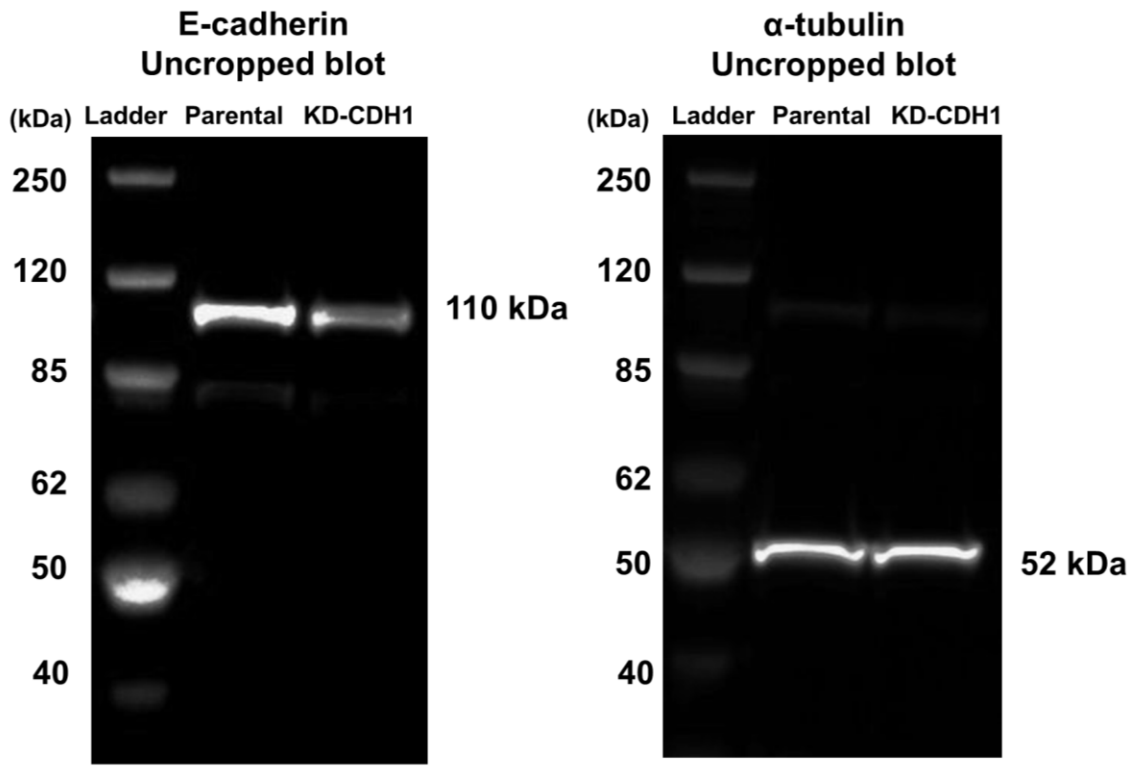


**Figure S4. Uncropped Western blots validating E-cadherin knockdown.** E-cadherin (left) and α-tubulin loading control (right) in parental and KD-CDH1 cells. Molecular weight markers in kDa at left of each panel.

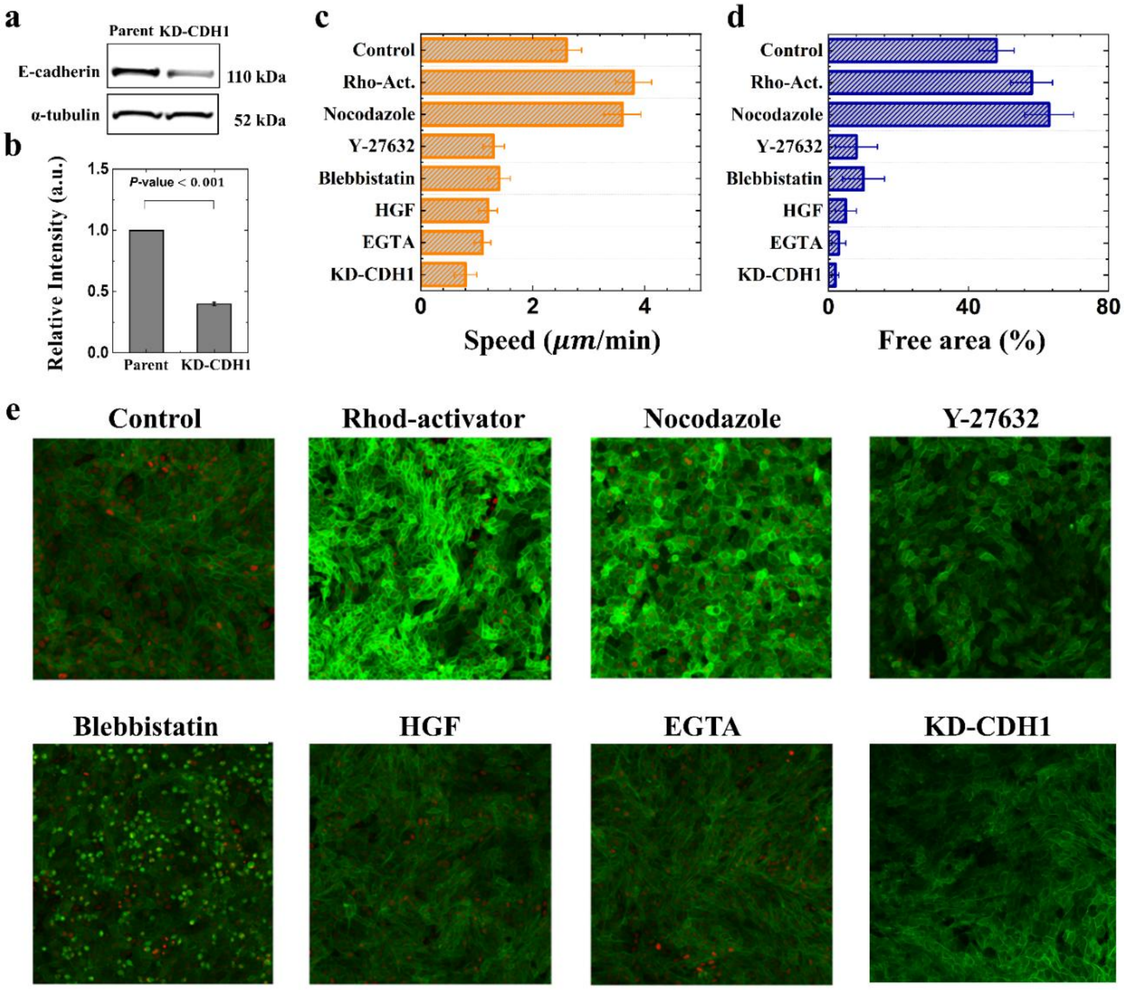


**Figure S5. Actomyosin contractility and E-cadherin-mediated adhesion are both required for collective reorganization.** (a,b) Western blot for E-cadherin and α-tubulin loading control (a) and densitometry (b) show E-cadherin reduced to 40 ± 1.5% of parental levels (N = 3). Two-sided Welch's two-sample t-test was used for statistic validation. N is the number of independent experiments. (c) Migration speed at 20 h under pharmacological or genetic perturbations as indicated. Error bars: SD, N = 3 per condition. (d) Free area fraction at 20 h for the same conditions. (e) Representative confocal fluorescence images for each perturbation.

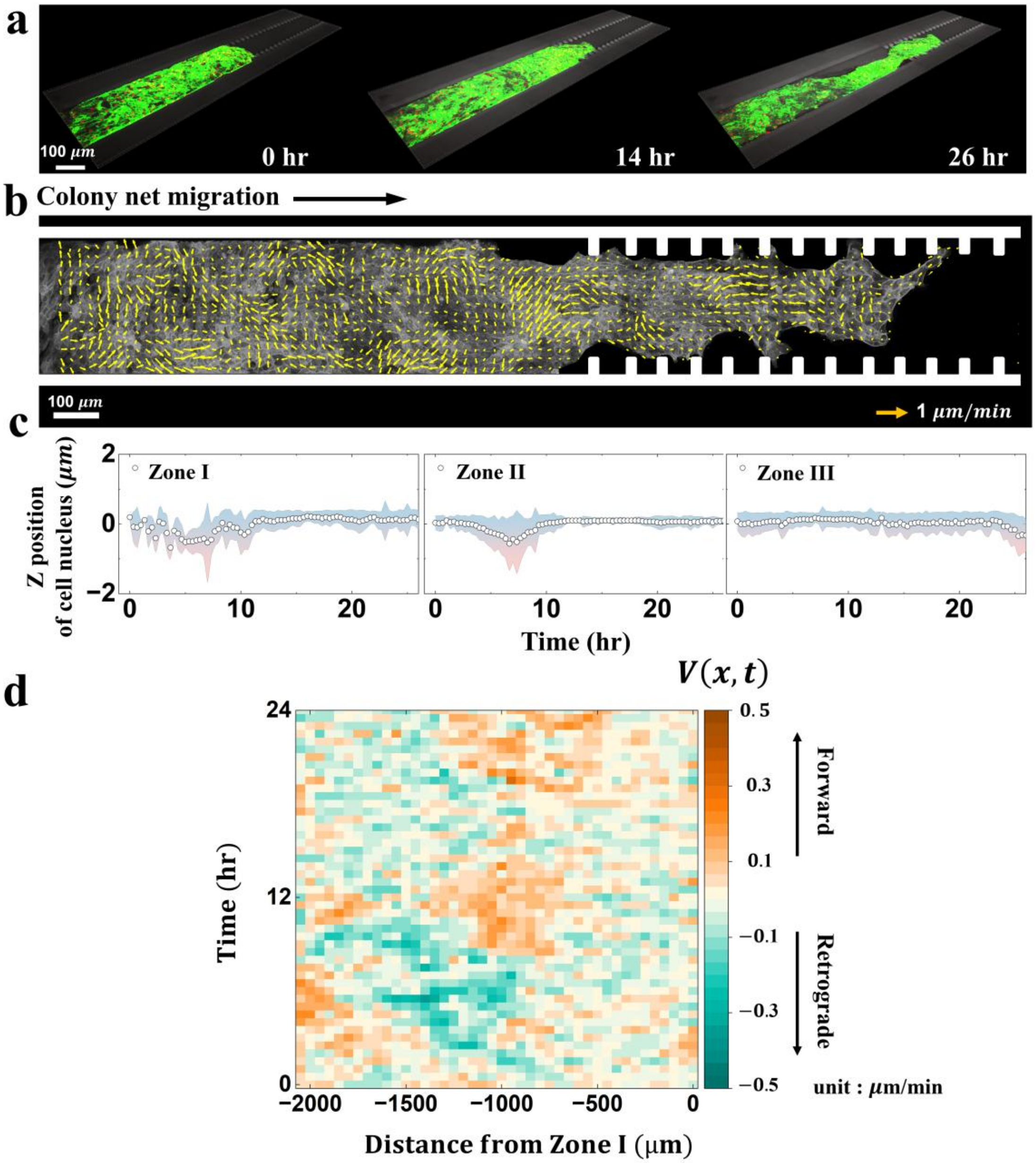


**Figure S6. Alternating flow persists at a serrated inlet**. (a) Compressed time-lapsed images of a migrating parental MDCK colony expressing lifeact-mEGFP (F-actin, green) at times 0, 14, and 26 h after cells contact the inlet. (b) PIV velocity field at 24 h. Arrows indicate local velocity direction and magnitude. (c) Mean z-position of 3D individual cell nuclei in Zones I, II and III; shaded region shows SD. Displacement stays within ±2 μm throughout, so the alternating flow in (d) reflects in-plane motion rather than cells moving out of the focal plane. (d) Kymograph of the axial velocity, $V(x,t)$.

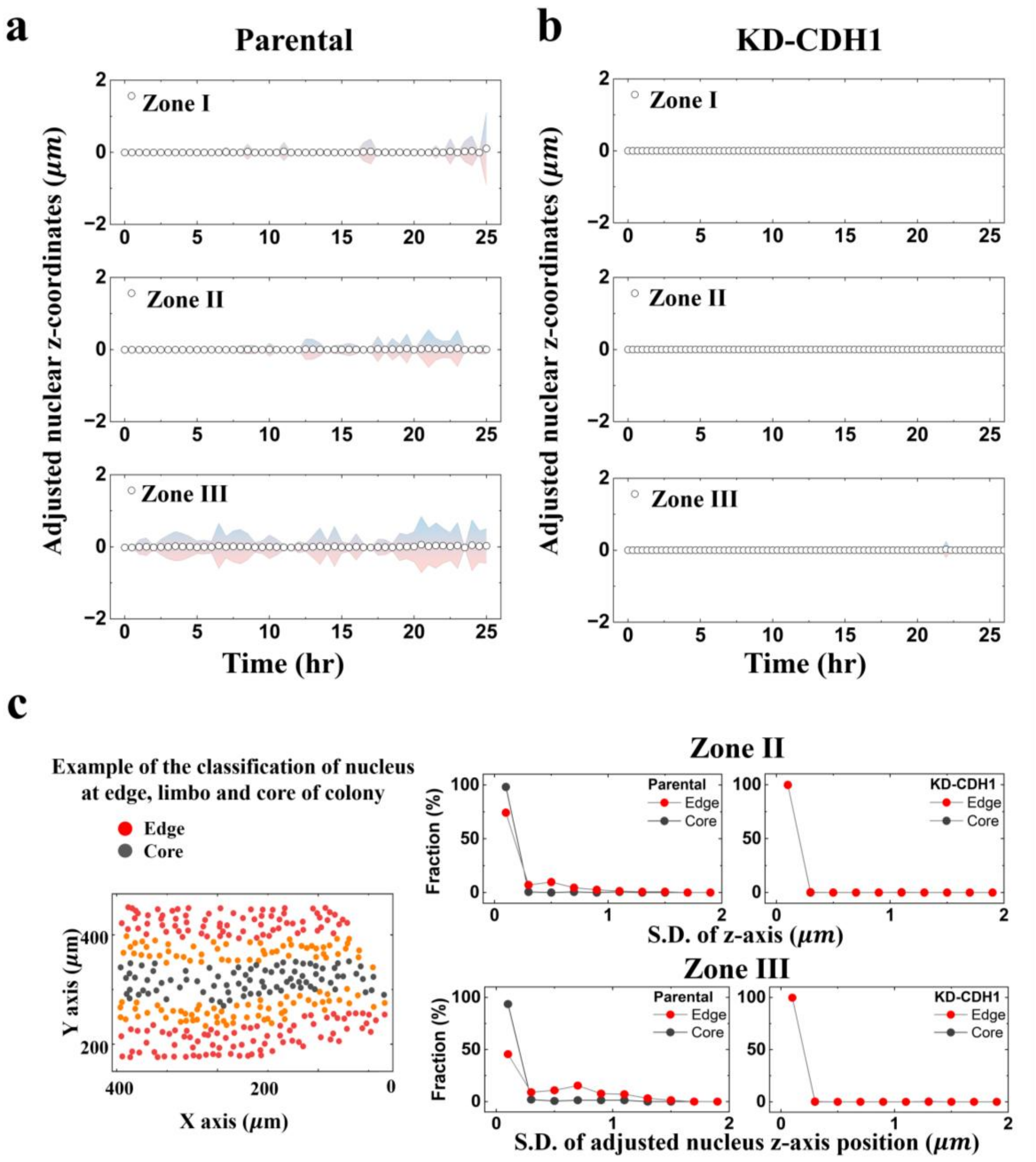


**Figure S7**. **Colony reorganization occurs in-plane, with nuclei displacement along the z axis below ±2 μm.** (a,b) The adjusted nuclear z-coordinates change over time in Zones I, II and III for parental (a) and KD-CDH1 cells (b). Displacement stays within ±2 μm throughout, so the alternating flow reported in Fig. 1 reflects in-plane motion rather than cells leaving the focal plane. (c) Nuclei near the colony boundary fluctuates more along z than nuclei in the interior in parental colonies, whereas edge and interior are indistinguishable for KD-CDH1 (N=3). N is the number of independent experiments. The example of classified nucleus depending on the distance fraction from the colony central line indicated as color (0-30%: core, dark gray; 70-100%: edge, red).

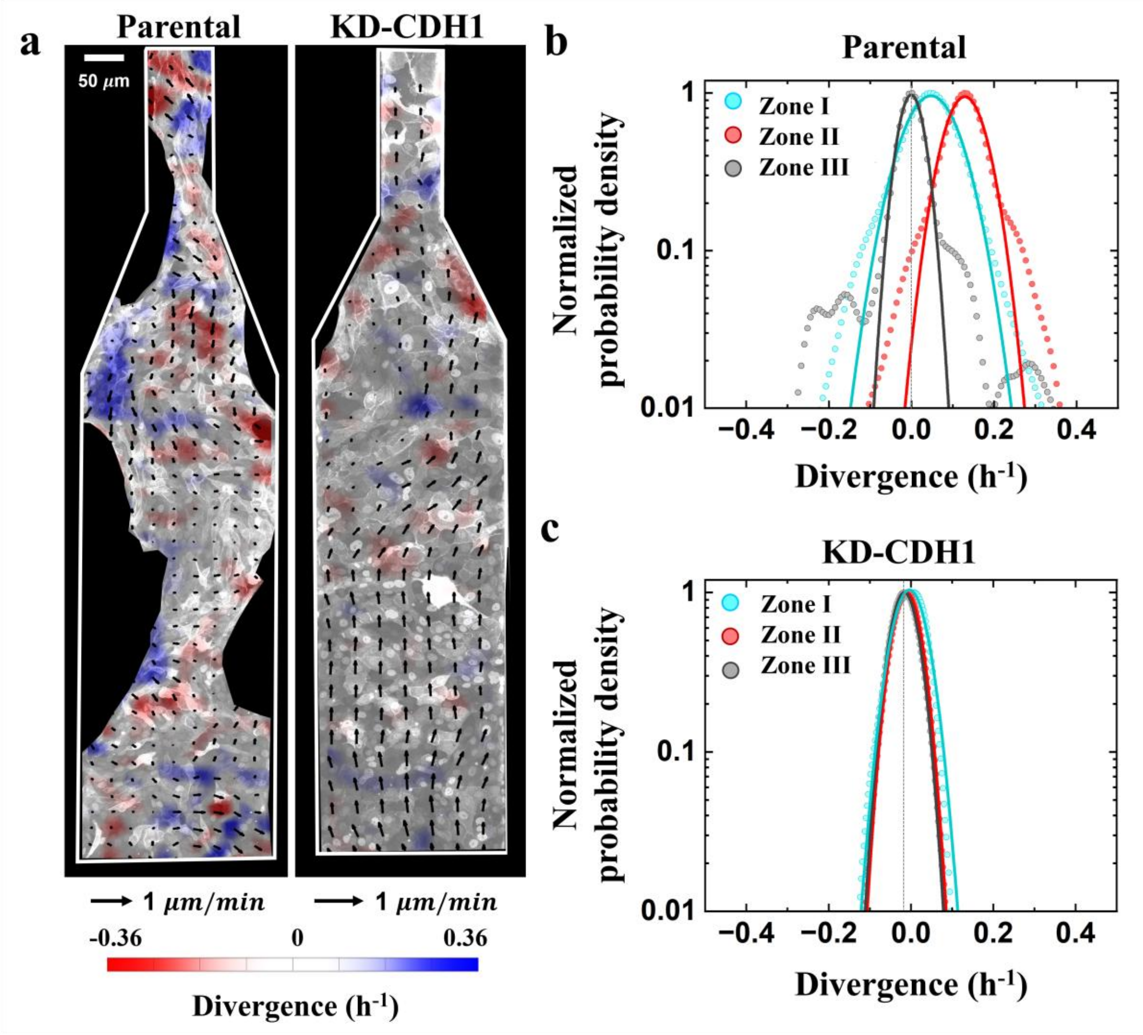


**Figure S8. Flow divergence is positive in Zone II of parental colonies and absent in KD-CDH1.** (a) Representative divergence maps for parental (left) and KD-CDH1 (right) colonies, inferred from PIV analysis. Positive values mark sources. Color scale in $h^{-1}$; scale bar, 50 $\mu$m. (b-c) Probability density of the regionally averaged divergence in Zones I, II, and III, for parental (b) and KD-CDH1 colonies, with Gaussian fits. Fits are in Table S3. N=3 independent experiments per genotype.

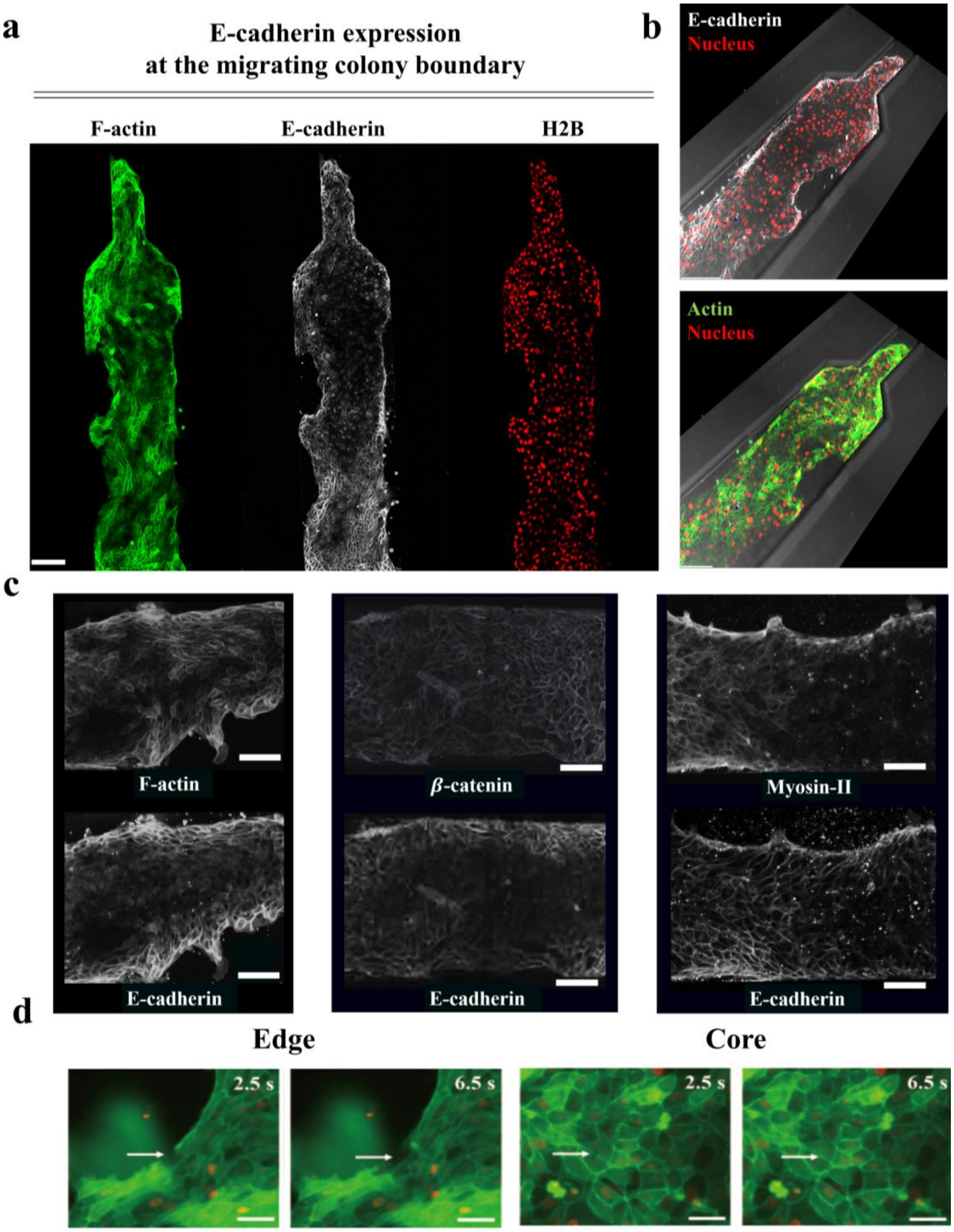


**Figure S9. E-cadherin, F-actin, and myosin-IIA co-enrich at the colony periphery; β-catenin does not.** (a-b) Confocal projection of the colony body viewed from above (a) and oblique view (b), showing E-cadherin accumulation at the edge (white) with F-actin (green) preserved in the interior. Nuclei are red. Scale bar is 100 $\mu$m. (c) Confocal images of two parental MDCK cell colonies at $t$ = 20 h, focusing between Zone II and Zone III regions. The samples are stained with F-actin and E-cadherin (left), β-catenin and E-cadherin (middle), and myosin-IIA and E-cadherin (right). The scale bar is 40 $\mu$m. (d) Laser ablation at the colony edge (left) and core (right); edge region is defined as within 40 $\mu$m of the colony boundary, core is the colony interior. Arrowheads show the ablation site. Scale bar: 15 $\mu$m.

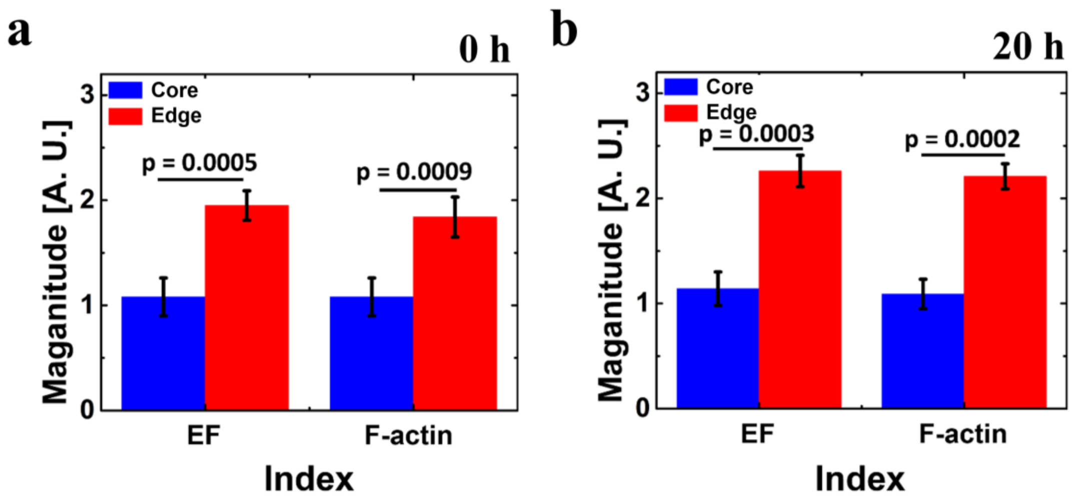


**Figure S10. Edge cells elongate and accumulate F-actin between 0 and 20 h.** (a,b) Cell elongation factor (EF) and F-actin expression, each normalized to the bulk colony edge, for edge and interior cells at t=0 h (a) and t=20 h (b). Edge is defined as within 40 $\mu m$ of the colony boundary.

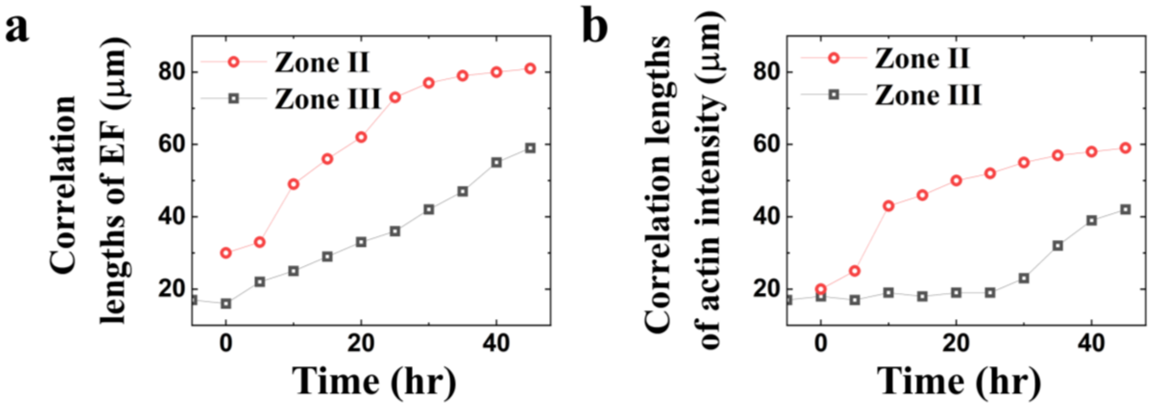


**Figure S11. Correlation lengths of cell elongation and F-actin intensity compared in Zones II and III.** (a) Cell elongation factor; (b) F-actin intensity.

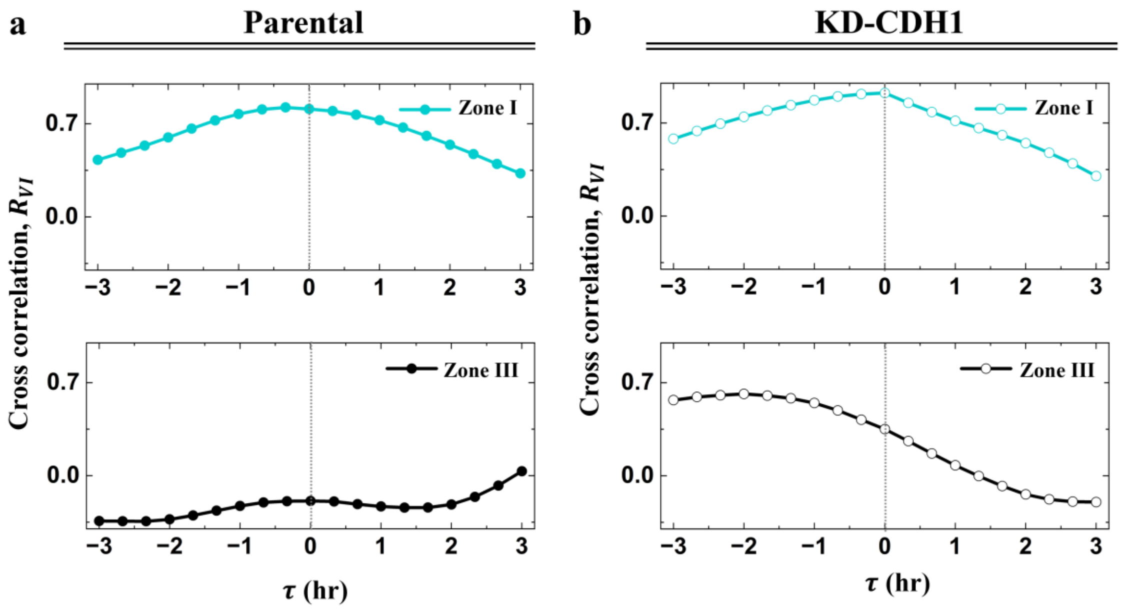


**Figure S12. Velocity and F-actin fluctuations are coupled without resolvable lag in Zones I and III.** (a, b) Cross-correlation $R_{VI}(\tau)$ between projected velocity and F-actin intensity in Zones I and III, for parental (a) and KD-CDH1 colonies (b). A peak at $\tau < 0$ indicates that velocity fluctuations precede F-actin fluctuations.

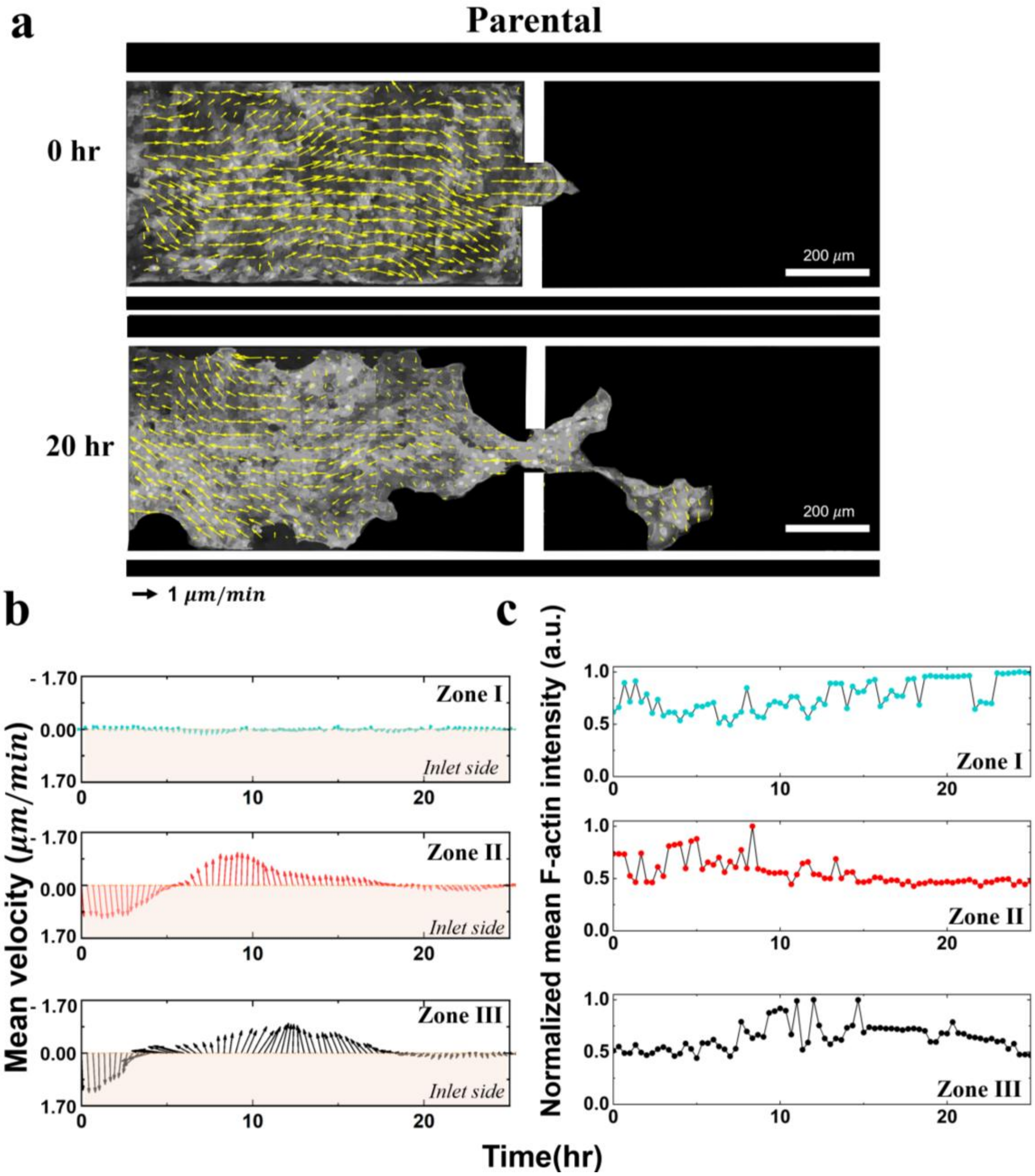


**Figure S13**. **Flow reversals and F-actin fluctuations persist at a constriction with entry angle 90° and remain E-cadherin dependent.** (a) PIV velocity fields at 0 h and 20 h for parental and KD-CDH1 entering the constriction. Scale bar, 200 μm. (b) Projected velocity in Zones I (top), II (middle) and III (bottom) regions, plotted against time. (c) Normalized f-actin intensity in the same regions against time. Mean ± SD; N=3, N is the number of independent experiments; time zero is first leading cells contact with the inlet.

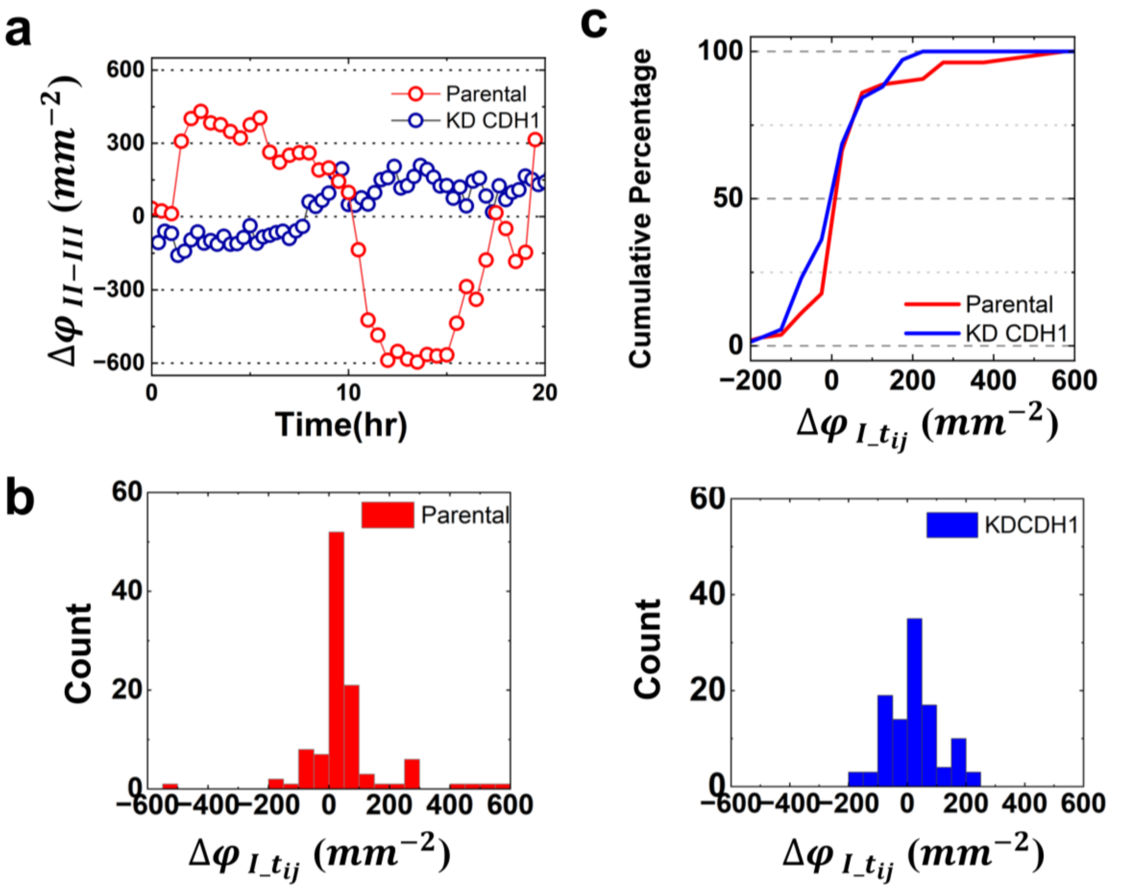


**Figure S14. Density difference between Zones II and III, and step-to-step density changes in Zone I.** (a) Difference in cell density between Zones II and III, for parental and KD-CDH1 colonies, plotted against time (N=3) (b) Change in the Zone I density between successive timepoints, for parental and KD-CDH1 colonies. (c) Cumulative distribution in Zone I for the two colony lines. Binned time is 20 min. (N=3, total data points = 109 for parental and KD-CDH1). N is the number of independent experiments.

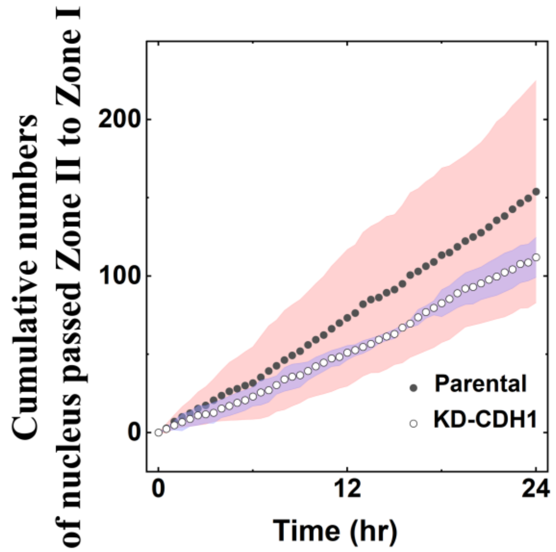


**Figure S15. Cumulative transit of nuclei from Zone II into Zone I, comparing parental and KD-CDH1 colonies.** Plotted against time. N=3, N is the number of independent experiments**.**

**Table S1. Recombinant DNA plasmids and shRNA oligonucleotide used modulate the protein expression of MDCK cell line.**

<table>
<tr><th colspan="2">Plasmid name</th><th>Source</th><th>Identifier</th><th>Purpose</th></tr>
<tr><td colspan="2">CMV-Lifeact-7- mEGFP</td><td>Addgene, USA</td><td>#54610</td><td>Localization of Lifeact-7 to Actin</td></tr>
<tr><td colspan="2">pcDNA3-CMV-H2B.1-mCherry</td><td>Addgene, USA</td><td>#20972</td><td>Histone H2B.1 fused to mCherry for chromatin visualization</td></tr>
<tr><td colspan="2">pLKO-TetOn-Puro</td><td>Addgene, USA</td><td>#21915</td><td>Lentivirus expression vector having tetracycline inducible shRNA expression</td></tr>
<tr><th colspan="2">Oligonucleotide</th><th>Source</th><th>Identifier</th><th>Purpose</th></tr>
<tr><td>Forward</td><td>5'-CCGGGCTCTCATTTCCGATTATATTCTCGAGTTCGAGAGTAAAGGCTAATATTTTTTG-3'</td><td rowspan="2">This paper</td><td rowspan="2">N.A.</td><td rowspan="2">Knock-down expression of CDH1 gene</td></tr>
<tr><td>Reverse</td><td>5'-AATTCAAAAAATATTAGCCTTTACTCTCGAACTCGAGAATATAATCGGAAATGAGAGC-3'</td></tr>
</table>

**Table S2. Pharmacological perturbations of actomyosin contractility and intercellular adhesion, and their effects on MDCK colony reorganization.**

| Chemical name | Supplier | Dose | Effect on confluent monolayers | Mechanism of action | Effect on upstream reorganization |
|---|---|---|---|---|---|
| Rho-activator | CN03-A (Cytoskeleton, USA) | 1 $\mu$g/ml | EF ↑ Actomyosin force ↑ | Activation of Rho-family GTPase | enhanced |
| Nocodazole | M1404 (Merck, USA) | 10 $\mu$M | Actomyosin force ↑ | Increase cell stiffness by activating GEF-H1/RhoA signaling and stabilization of junctions | enhanced |
| Y-27632 | SCM075 (Merck, USA) | 20 $\mu$M | EF ↓ Actomyosin force ↓ | Inhibit the activity of ROCK | suppressed |
| Blebbistatin | B0560 (Merck, USA) | 20 $\mu$M | Actomyosin force ↓ Extrusion ↑ | Inhibit the activity of myosin II | suppressed |
| HGF | H9661 (Merck, USA) | 20 ng/ml | Speed ↑ Confluence ↓ | Weaken cell-to-cell interaction by MAPK/Egr-1-mediated upregulation of Snail | suppressed |
| EGTA | 03777 (Merck, USA) | 200 $\mu$M | Speed ↑ Confluence ↓ | Weaken cell-to-cell interaction as a calcium specific chelator. Perturbing calcium dependent cell-cell adherence development | suppressed |

**Table S3. Gaussian fits of the distribution of regional flow divergence, by zone and cell line.** Here σ denotes the width of the fitted; it reports the spread of the underlying distribution and should not be read as an error bar.

| Zone | Center peak ($x_c$) ($h^{-1}$) | | Standard deviation ($\sigma$) ($h^{-1}$) | |
|---|---|---|---|---|
| | **Parental** | **KD-CDH1** | **Parental** | **KD-CDH1** |
| **I** | 0.05 | < 0.01 | 0.06 | 0.04 |
| **II** | 0.13 | -0.01 | 0.05 | 0.03 |
| **III** | < 0.01 | -0.02 | 0.03 | 0.03 |

**Table S4. The cross-correlation at lag time $\tau$ = 0 hr of axial velocity between adjacent zones, parental and KD-CDH1. (N=3 independent batches).**

| Zone | Parental | KD-CDH1 |
|---|---|---|
| **Zone I vs Zone II** | 0.02 $\pm$ 0.30 | 0.27 $\pm$ 0.25 |
| **Zone II vs Zone III** | 0.67 $\pm$ 0.13 | 0.32 $\pm$ 0.44 |

**Legends for Supplementary Movies**

**Supplementary Movie 1. Parental colony entering the tapered inlet from the wide channel.** A typical confocal video of a MDCK-H2B-mCherry & Lifeact-mEGFP cell colony migrating into a narrow inlet from a wide channel. The green and red signal intensities are proportional to the local concentrations of F-actin and H2B, respectively. Frame interval, 20 min; total duration, 25 h; playback, 6fps.

**Supplementary Movie 2. Parental colony entering a curved inlet.** As Movie S1, in a curved channel. Frame interval, 20 min; total duration, 20 h; playback, 6fps.

**Supplementary Movie 3. KD-CDH1 colony entering the tapered inlet.** As Movie S1, for a MDCK-KD-CDH1 colony. Frame interval, 20 min; total duration, 24 h; playback, 6fps.

**Supplementary Movie 4. Parental colony entering a fibronectin-coated tapered inlet.** As Movie S1, with the channel coated with fibronectin at 10 μg/mL. Frame interval, 20 min; total duration, 35 h; playback, 6fps.

**Supplementary Movie 5. Flow at the colony boundary in the wide channel.** PIV velocity fields overlaid on the F-actin signal (grey), shown for three conditions in sequence: parental on uncoated PDMS, KD-CDH1 on uncoated PDMS, parental on fibronectin-coated PDMS. Acquisition and playback as in Movie S1.
